# Neuroanatomical and behavioral characterization of corticotropin releasing factor-expressing lateral Habenula neurons in mice

**DOI:** 10.1101/2025.04.29.651264

**Authors:** William J. Flerlage, Shawn C. Gouty, Emily H. Thomas, Sarah C. Simmons, Mumeko C. Tsuda, Oana L. Rujan, Regina C. Armstrong, Catherine M. Davis, Laxmi Iyer, Emily Petrus, Brian M. Cox, T. John Wu, Fereshteh S. Nugent

## Abstract

The lateral habenula (LHb) is a critical hub for stress-related behaviors, yet the sources of its corticotropin-releasing factor (CRF) inputs remain poorly defined. Using high-resolution imaging, RNAscope, and viral tracing, we identified a novel, intrinsic population of CRF-expressing LHb neurons (LHb^CRF^). These neurons are primarily VGLUT2+, though a rostral subpopulation co-expresses GAD2. While chemogenetic activation of LHb^CRF^ neurons did not impact place preference or anxiety-like behaviors, it selectively biased defensive strategies toward passive action-locking during the Visual Looming Shadow Test (VLST). Notably, this activation prolonged escape latencies in males and post-escape shelter stays in females. Electrophysiological and optogenetic characterization revealed significant sexual dimorphism: male LHb^CRF^ neurons are more numerous and intrinsically excitable, whereas female LHb^CRF^ neurons exhibit stronger local excitatory connectivity. These findings establish LHb^CRF^ neurons as a sexually dimorphic circuit component that could modulate sex-specific defensive strategies under threat via divergent cellular and synaptic mechanisms between the sexes.

## Introduction

The lateral habenula (LHb) is an evolutionarily conserved brain region crucial for behavioral selection, in response to stressors and rewards; thereby guiding goal-directed motivated behavior^1–3^. Ongoing research continues to unveil nuanced roles for the LHb in both normal behavior and the pathophysiology of mental illness. Aversive stimuli as well as the omission of expected reward generally activate the LHb, and its hyperactivity is implicated in anhedonic and negative states associated with depression, anxiety, and drug addiction. Conversely, reward-related stimuli inhibit LHb activity which generates positive motivational states. These findings highlight LHb’s therapeutic potential, as evidenced by the anti-depressant effects of deep brain stimulation of the LHb and ketamine’s NMDA receptor antagonism of LHb neurons ^1–5^.

The majority of LHb neurons are glutamatergic, and express markers such as the vesicular glutamate transporter 2 (VGLUT2) ^6^. However, a small subset also expresses classical GABAergic markers, including glutamic acid decarboxylase 1 (GAD1), GAD2, the GABA transporter 1 (GAT-1), and the vesicular GABA transporter (VGAT), suggesting the presence of a functional intrinsic inhibitory circuit within the LHb^7–10^.

Notably, some LHb neurons co-express both glutamatergic and GABAergic markers. For example, parvalbumin-expressing LHb neurons (LHb^PV^) comprise three distinct subpopulations: a majority that are VGLUT2+ and glutamatergic, a subset that expresses only GABAergic markers, and a subpopulation that co-expresses both glutamatergic and GABAergic markers (e.g., VGLUT2 and GAD2)^7^. Most LHb glutamatergic outputs directly and indirectly modulate key monoaminergic systems, primarily the dopamine and serotonin pathways that encode behavioral avoidance and reward prediction errors. Mechanistically, this involves suppressing dopamine release from the ventral tegmental area (VTA) and serotonin release from the raphe nuclei by exciting GABAergic neurons, particularly those in the rostromedial tegmental area (RMTg), thus shifting behavior away from reward-seeking^2,12–14^. Consequently, LHb is increasingly recognized as a critical structure in the neural circuits that govern mood, motivation, and decision-making by processing a diverse array of appetitive and aversive signals that are crucial for adaptive behavior, allowing organisms to learn from both positive and negative experiences^5^.

LHb neuronal activity is regulated by glutamatergic, GABAergic, and glutamate/GABA co-releasing synaptic inputs from various regions in the forebrain and midbrain such as the medial prefrontal cortex^15,16^, entopeduncular nucleus/globus pallidus^17,18^, lateral preoptic area (LPO) ^29^, ventral pallidum^19^, lateral hypothalamus (LH/LHA) ^20^, ventromedial hypothalamus^16,21,22^ , septum^23^, central amygdala (CeA) ^21^, and bed nucleus of the stria terminalis (BNST)^21^. It also receives reciprocal inputs from the VTA^24^, laterodorsal tegmental nucleus^25^, periaqueductal gray (PAG)^15,16^, and locus coeruleus^1,15^ forming complex feedback loops that modulate activity. Given the widespread expression of various G protein-coupled receptors on LHb neurons and their afferents, LHb neuronal activity is subject to neuromodulatory actions of both fast and slow synaptic transmission from glutamate and GABA, as well as other neurotransmitters and neuromodulators including endocannabinoids^26^, acetylcholine^27^, serotonin^28–30^, norepinephrine^31^ and several co-released neuropeptides such as orexin^8^, neuropeptide Y^32,33^, dynorphin^16,34^ and corticotropin-releasing factor (CRF)^35–38^. These diverse glutamatergic, GABAergic, and neuromodulatory signals provide the LHb with information related to motivation, reward, aversion, and stress.

Specifically regarding CRF neuromodulatory actions, hypothalamic (paraventricular nucleus of hypothalamus, PVH) and extra-hypothalamic CRF systems mediate stress responses, and their dysfunction is implicated in stress-related disorders. CRF’s central action through its receptors CRFR1/2 (both G protein-coupled receptors) is cell type-, region- and pathway-specific and depends on prior stress exposure^39^. Additionally, sex differences in CRF signaling may contribute to female vulnerability to anxiety and depression^40^. Our previous studies demonstrated that the rodent LHb is highly responsive to CRF^35,36,38^, increasing LHb neuronal excitability via CRFR1-PKA-dependent suppression of different potassium currents (small conductance potassium SK channels in rat LHb^38^; M-type potassium channels in mouse LHb^35^). In rat LHb, CRF reduces presynaptic GABA release via retrograde endocannabinoid signaling, without changing glutamatergic activity, whereas in mouse LHb, CRF attenuates both GABAergic and glutamatergic transmission^35,38^. We also showed that early life stress and brain trauma augment endogenous LHb CRF-CRFR1 signaling, contributing to LHb hyperexcitability^36,38^.

Building on these findings, here we investigated the source of CRF inputs to the LHb which remained unclear. Using viral-based Cre-mediated strategies in CRF transgenic mouse lines, we mapped LHb-projecting CRF neurons and discovered a previously uncharacterized neural subpopulation of LHb that expresses CRF (LHb^CRF^ neurons). We then traced the major downstream targets of LHb^CRF^ axonal projections, revealing a complex architecture of both local intra-habenular microcircuits and widespread long- range projections. Our mapping identified LHb^CRF^ terminals across multiple brain regions, including diencephalic regions such as the LH/LHA and zona incerta (ZI), as well as critical midbrain and hindbrain centers involved in defense and motivation, such as the PAG, superior colliculus (SC), VTA, and RMTg. For the initial characterization of behavioral relevance of these neurons, we employed a chemogenetic gain-of-function strategy using Gq-DREADDs. This approach was specifically chosen over inhibition because chemogenetic silencing of major LHb projection targets such as the VTA and RMTg often fail to modulate baseline behaviors like locomotion or anxiety unless the animal is specifically challenged by a stressor^15^. Furthermore, given that the chemogenetic activation of other distinct LHb subpopulations, such as Pituitary Adenylate Cyclase-Activating Polypeptide (PACAP)-expressing neurons, has been shown to paradoxically produce rewarding effects and decrease fear^41^, we sought to determine the specific valence and behavioral impact of the LHb^CRF^ population upon their activation. We utilized a battery of assays including the Open Field Test (OFT), Elevated Zero Maze (EZM), and Light-Dark Task (LDT) as critical controls to ensure that chemogenetic activation of LHb^CRF^ neurons did not simply induce generalized anxiety or motor deficits. To test the role of these neurons in motivated and defensive states, we employed Conditioned Place Preference (CPP) and the Visual Looming Shadow Test (VLST). Specifically, the VLST was chosen for the assessment of innate defensive behaviors, as hypothalamic CRF neurons and LHb neurons have been shown to play a significant role in the modulation of this behavior ^42–45^. Our results reveal that LHb^CRF^ neurons may play a specific role in innate threat-evoked defensive behaviors, characterized by increased frequency and duration of action-locking behaviors and sex-dependent differences in escape latencies and post-escape safety-seeking upon chemogenetic activation. We also identified significant sexual dimorphism in the population size, basal excitability, and local excitatory connectivity of LHb^CRF^ neurons. These findings suggest that while both sexes exhibit an increased adoption of action-locking behaviors upon LHb^CRF^ circuit activation, they may do so through potentially divergent cellular and synaptic mechanisms. Specifically, the higher absolute number and increased intrinsic excitability of LHb^CRF^ neurons in males, contrasted with the significantly stronger local synaptic gain observed in females, may contribute to the observed sex-dependent variations in escape latency and post-escape safety-seeking behavior.

## Results

In the present study, we used CRH-Cre::Ai14 or CRH-Cre::Ai6 reporter lines in conjunction with the standard CRH-Cre line to facilitate the specific identification and visualization of CRF+ neurons for our RNAscope, electrophysiology, local circuit mapping as well as chemogenetic and optogenetic experiments. It is important to note that all three lines utilize the same Cre-driver to target the CRF population; the reporter lines (Ai14/Ai6) simply offer the technical advantage of endogenous fluorophore expression (tdTomato or ZsGreen) upon Cre-mediated recombination. Extensive literature has validated that these Cre-dependent reporter strains reliably recapitulate the expression patterns of the underlying Cre-driver line without altering the physiological or behavioral characteristics of the target neuronal population ^46,47^.

Specifically, the CRH-Cre line has been widely demonstrated to target CRF neurons with high specificity and efficiency across various brain regions, and the addition of the Ai reporter alleles does not introduce confounding variables in behavioral responses^45,48–52^.

### A subpopulation of CRF projections to the LHb originates within the LHb itself

To identify CRF projections to the LHb, we used Cre-dependent retrograde viral tracing by bilaterally injecting retrograde adeno-associated virus (rgAAV-DIO-YFP) into the LHb of CRH-Cre male and female mice (n=3, Figure 1 and Supplementary Figure 1). This revealed an intrinsic subpopulation of LHb neurons expressing CRF (LHb^CRF^).

**Figure 1:**
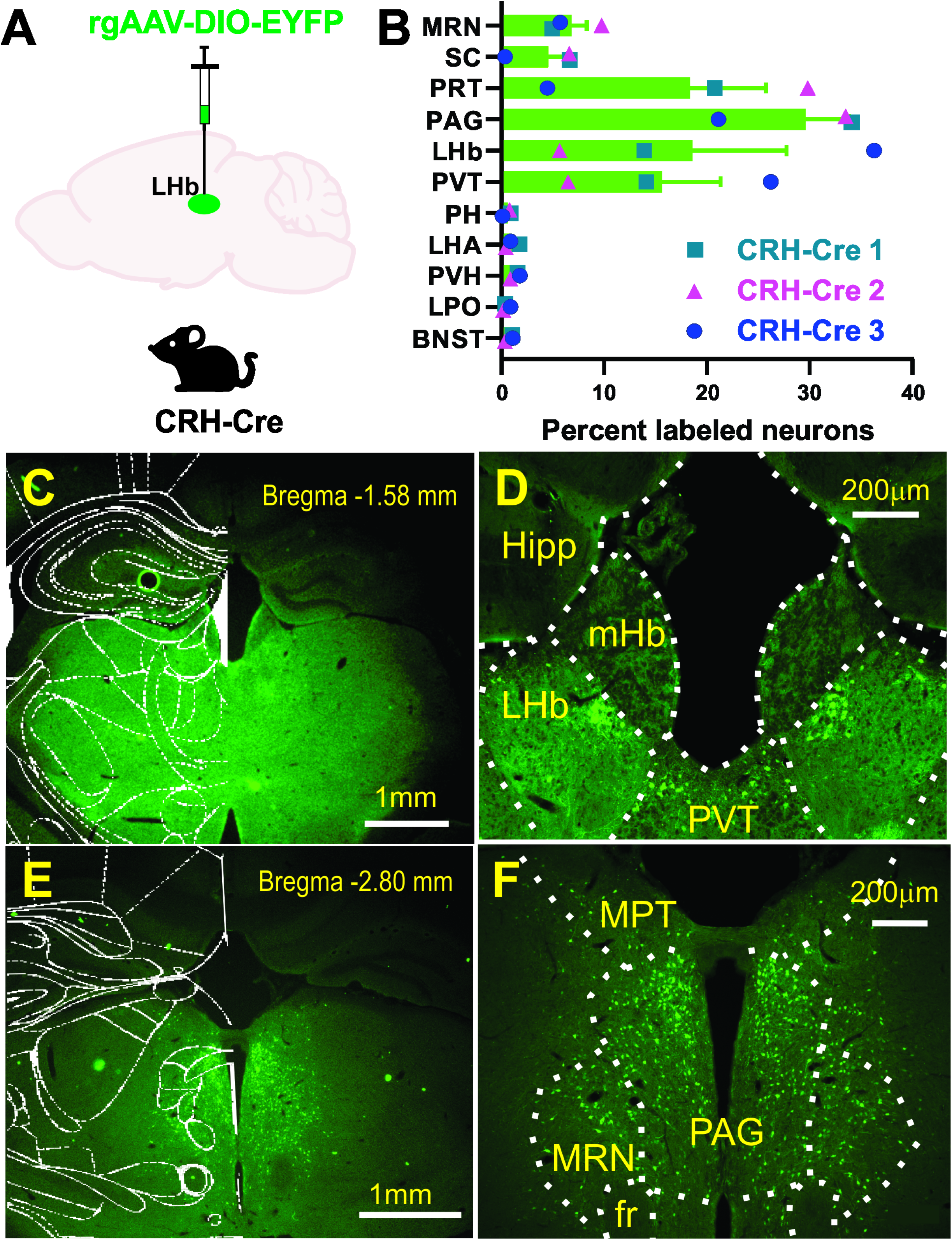
Retrograde Identification of CRF-Expressing Projections to the Lateral Habenula (LHb) **A)** Schematic of the viral strategy using a Cre-dependent retrograde vector (rgAAV-DIO-EYFP) injected into the LHb of CRH-Cre mice. This approach selectively labels the cell bodies of CRF-expressing neurons that provide direct input to the LHb. **B)** Quantitative distribution of retrogradely labeled cell bodies across identified brain regions. Data are presented as the percentage of total labeled neurons per animal (n=3 mice: 2 males, 1 female). Individual symbols (circles, triangles, squares) represent data from single subjects (CRH-Cre 1–3); bars indicate the mean. High densities of labeled cells were observed in the pretectal area (PRT), periaqueductal gray (PAG), LHb, and paraventricular thalamic nucleus (PVT). Total cells analyzed per subject: CRH-Cre 1 (n=2211), CRH-Cre 2 (n=3309), and CRH-Cre 3 (n=1750). **C, E:** Low-magnification coronal sections with mouse atlas overlays at the indicated Bregma levels (−1.58 mm and −2.80 mm), showing the spatial distribution of EYFP+ neurons. **D, F:** High-magnification images of key regions of interest (ROIs). **D** highlights labeling in the LHb and PVT; **F** illustrates labeling in the medial pretectal nucleus (MPT, part of the PRT), PAG, and midbrain reticular nucleus (MRN). Supplementary Figure 1 contains representative images of the remaining brain regions identified as containing CRF+ neurons projecting to the LHb. Abbreviations: *BNST*, bed nucleus of the stria terminalis; *LPO*, lateral preoptic area; *LHA*, lateral hypothalamus; *PH*, posterior hypothalamus; *PVH*, paraventricular hypothalamic nucleus; *SC*, superior colliculus; *fr*, fasciculus retroflexus. Scale bars: C, E = 1mm; D, F = 200μm.

Beyond this local population, our mapping identified several prominent long-range CRF-expressing inputs:

- Primary Afferents: High densities of retrogradely labeled CRF+ cell bodies were observed in the pretectal area (PRT), the paraventricular nucleus of the thalamus (PVT), and the PAG.
- Secondary Midbrain Inputs: Substantial labeling was also identified in the midbrain reticular nucleus (MRN) and SC.
- Hypothalamic and Limbic Contributions: Smaller but consistent populations of CRF+ neurons were detected in the PVH, posterior hypothalamus (PH), LH/LHA, BNST, and LPO.

To determine if LHb^CRF^ neurons constitute a subclass of LHb^PV^ neurons, we also performed anti-parvalbumin immunostaining on retrogradely labeled LHb^CRF^ neurons. The absence of detectable parvalbumin expression in LHb^CRF^ neurons indicates that these may constitute distinct subpopulations, separate from LHb^PV^ neurons. (Supplementary Figure 2).

### LHb^CRF^ Neurons express VGLUT2 and GAD2 with a minority expressing vGAT mRNA

To characterize the neurochemical identity of LHb^CRF^ neurons, we performed a comprehensive, large-scale RNAscope experiment utilizing Zeiss AxioScan imaging and QuPath automated analysis (Supplementary Figure 3), across the rostrocaudal axis spanning five AP levels. While we initially attempted to utilize a Crh probe, technical limitations regarding non-specific binding led us to utilize CRH-Cre::Ai14 mice, where the tdTomato reporter identifies CRF+ neurons. Our data (Figures 2–3 and Supplementary Figures 3–7) suggest that LHb^CRF^ neurons are primarily concentrated in the rostral portion of the LHb (Bregma −1.06 to −1.34 mm, Figures 2-3 and Supplementary Figures 4–7). Within these levels, the population exhibits a distinct spatial organization: a dense cluster is situated in the medial LHb (LHbM), bordering the medial habenula, while a significant subpopulation extends into the lateral LHb (LHbL). As the axis moves caudally (Bregma −1.58 mm, Supplementary Figures 6-7), the population density decreases significantly. To ensure our quantification was not confounded by potential structural dimorphism, we analyzed the total surface area of the LHb across all five AP levels using the Allen Brain Atlas and Paxinos templates. We identified a total of 1,388 CRF+ cells across 13 mice (n = 833 in 7 males; n = 555 in 6 females). We found no significant differences in LHb surface area between sexes (Supplementary Figure 3; F(1, 55) = 1.924, p = 0.17, 2-way ANOVA), allowing for an unbiased comparison of cell counts. Our results show that while the topographical distribution is identical between sexes, male mice possess a significantly higher absolute number of LHb^CRF^ neurons than females (Figure 4; Student’s t-test, p < 0.05). We further characterized these cells by their transcriptional profile, confirming that 100% express VGLUT2. Approximately 17–18% co-express GAD2, while a negligible minority (2–5%, primarily in males) co-express vGAT and VGLUT2 (Figure 4; Chi-Square test, p < 0.01). These findings identify LHb^CRF^ neurons as a predominantly VGLUT2+ subpopulation with some co-expressing GAD2+ and a significant sex-dependent variations in total population size.

**Figure 2:**
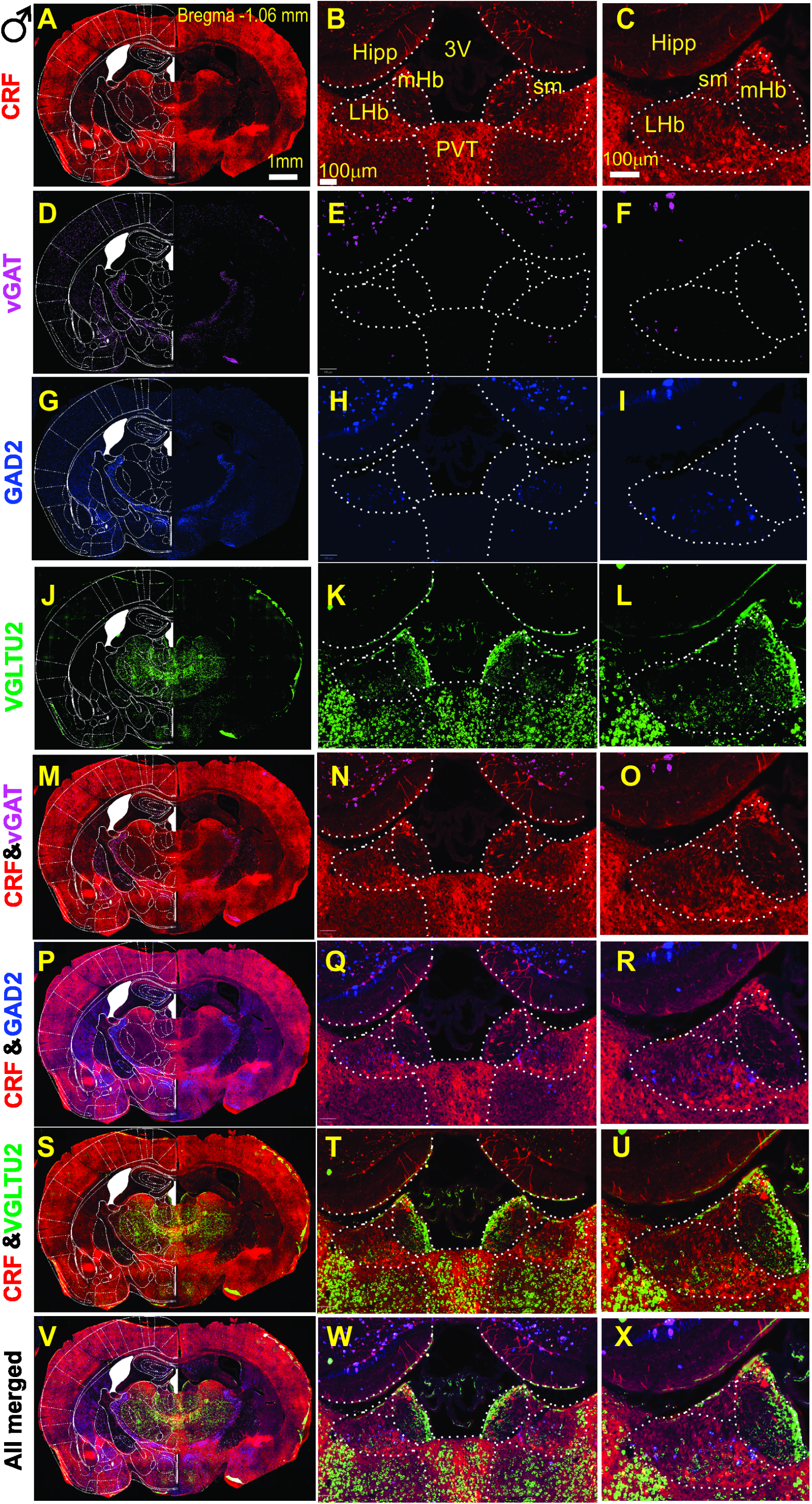
Molecular Characterization of CRF+ Neurons in the male LHb at Bregma −1.06mm. Multiplexed RNAscope *in situ* hybridization was utilized to identify the neurotransmitter phenotype of CRF+ (tdTomato-labeled) neurons in male CRH-Cre::Ai14 mice at Bregma −1.06 mm. (**A**, **D**, **G**, **J**): Low magnification hemicoronal views integrated with mouse atlas overlays, showing regional expression patterns of *Crh* (tdTomato), *Slc32a1* (vGAT), *Gad2* (GAD2), and *Slc17a6* (vGLUT2). (**B**–**L**): High-magnification images corresponding to panels A, D, G, and J, highlighting bilateral expression within the lateral habenula (LHb) and providing high-resolution details of the left LHb. (**M**–**U**): Dual-channel merged images demonstrating the colocalization (overlay) of CRF with inhibitory markers (*vGAT*, *GAD2*) and the excitatory marker (*vGLUT2*). (**V**–**X**): All-channel merged images illustrating the simultaneous colocalization of *vGAT*, *GAD2*, and *vGLUT2* within LHb CRF+ neurons. Abbreviations: *LHb*, lateral habenula; *mHb*, medial habenula; *Hipp*, hippocampus; *sm*, stria medullaris; *3V*, third ventricle; *PVT*, paraventricular nucleus of the thalamus. Scale bars: A = 1mm; B, C = 100 μm.

**Figure 3:**
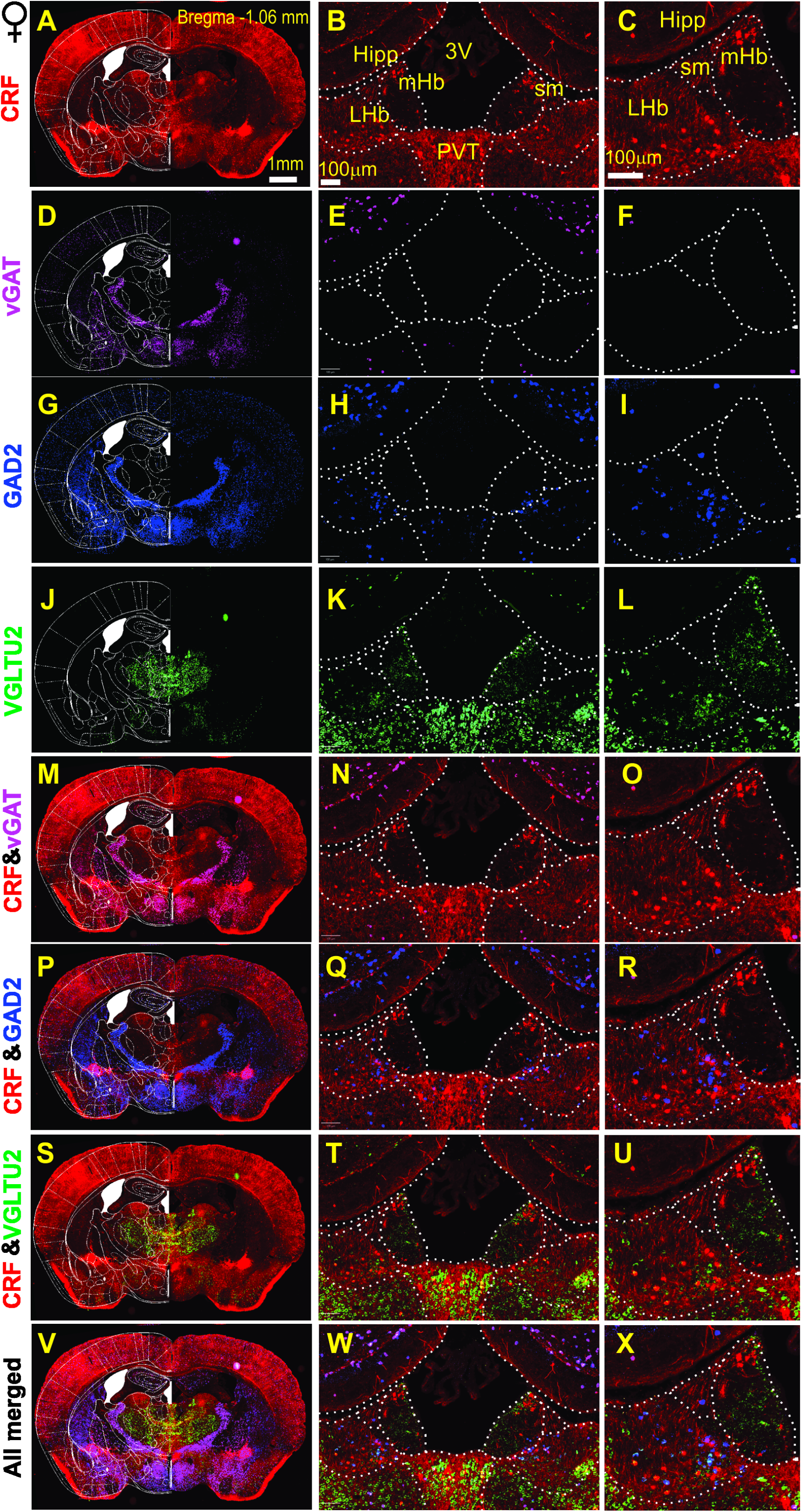
Molecular Characterization of CRF+ Neurons in the female LHb at Bregma −1.06mm. Multiplexed RNAscope *in situ* hybridization was utilized to identify the neurotransmitter phenotype of CRF+ (tdTomato-labeled) neurons in female CRH-Cre::Ai14 mice at Bregma −1.06 mm. (**A**, **D**, **G**, **J**): Low magnification hemicoronal views integrated with mouse atlas overlays, showing regional expression patterns of *Crh* (tdTomato), *Slc32a1* (vGAT), *Gad2* (GAD2), and *Slc17a6* (vGLUT2). (**B**–**L**): High-magnification images corresponding to panels A, D, G, and J, highlighting bilateral expression within the lateral habenula (LHb) and providing high-resolution details of the left LHb. (**M**–**U**): Dual-channel merged images demonstrating the colocalization (overlay) of CRF with inhibitory markers (*vGAT*, *GAD2*) and the excitatory marker (*vGLUT2*). (**V**–**X**): All-channel merged images illustrating the simultaneous colocalization of *vGAT*, *GAD2*, and *vGLUT2* within LHb CRF+ neurons. Abbreviations: *LHb*, lateral habenula; *mHb*, medial habenula; *Hipp*, hippocampus; *sm*, stria medullaris; *3V*, third ventricle; *PVT*, paraventricular nucleus of the thalamus. Scale bars: A = 1mm; B, C = 100 μm.

**Figure 4:**
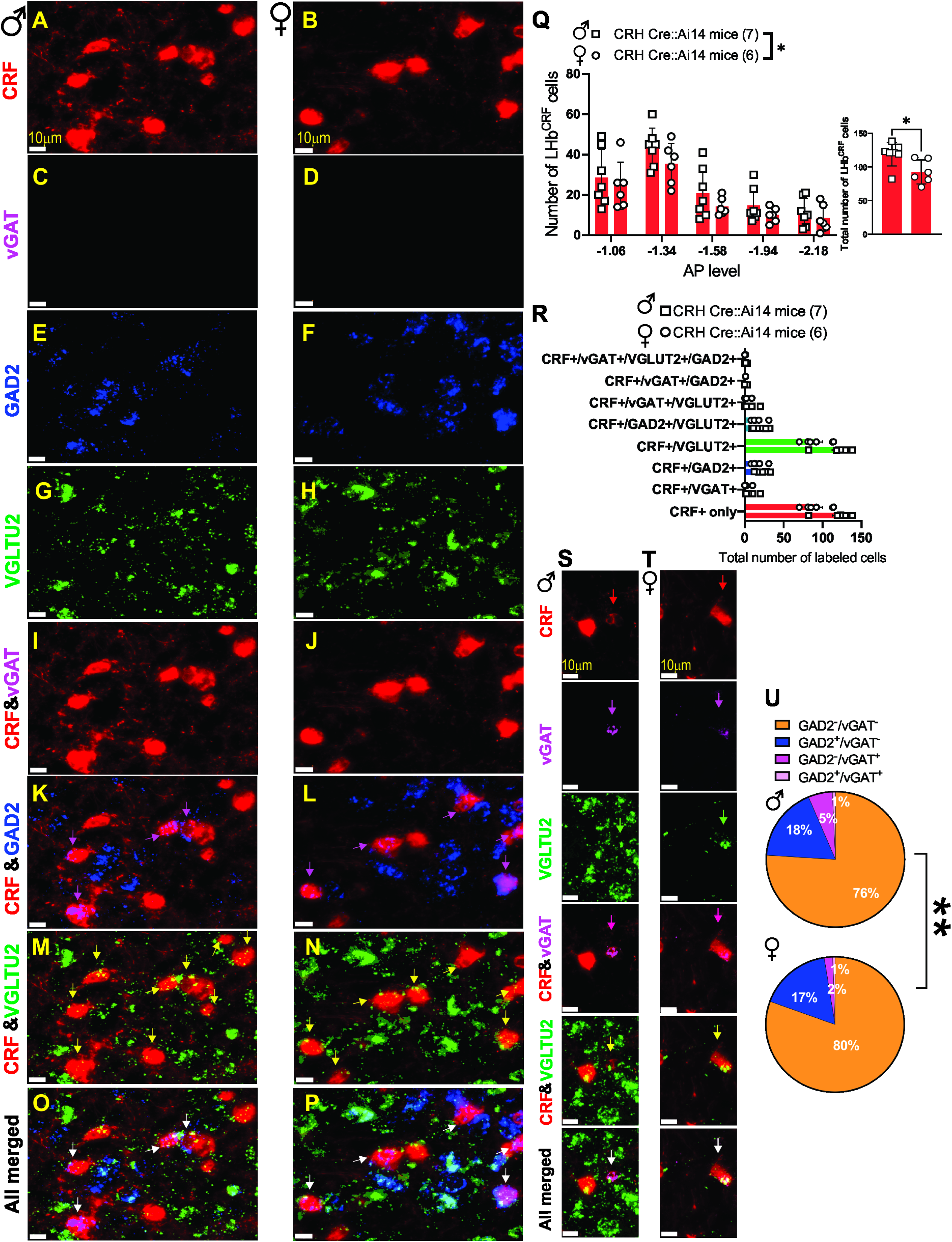
LHb^CRF^ neurons are predominantly glutamatergic and exhibit sex-dependent distribution. (A–P) High-resolution representative images from male (left columns) and female (right columns) mice showing the distribution of *Slc32a1* (vGAT), *Gad2* (GAD2), and *Slc17a6* (VGLUT2) mRNA. Given the low prevalence of the *vGAT+/CRF+* subpopulation (quantified in R and U), panels A–P represent the predominant neurochemical phenotypes: CRF+ neurons expressing vGLUT2 alone, GAD2 alone, or the co-expression of both markers. **(K, L):** Magenta arrows indicate CRF+ neurons co-expressing GAD2. **(M, N):** Yellow arrows indicate CRF+ neurons co-expressing VGLUT2. **(O, P):** White arrows highlight representative neurons showing triple colocalization of CRF, vGLUT2, and GAD2. **(Q)** Analysis of the number of CRF+ neurons identified per section across five anterior-posterior (AP) levels (−1.06 to −2.18 mm). The inset bar graph represents the total summation of CRF+ neurons across all analyzed levels. Male mice (squares, n=7) exhibited a significantly higher number of total LHb CRF+ neurons compared to females (circles, n=6). **(R)** Bar graph illustrating the frequency of total CRF+ neurons expressing various combinations of GABAergic and glutamatergic markers. Notably, 100% of identified CRF+ neurons co-express VGLUT2. While the vast majority are purely glutamatergic, small subpopulations exhibit dual-phenotypes, co-expressing either GAD2 or vGAT in addition to vGLUT2. No CRF+ neurons were found to be exclusively GABAergic. **(S–T)** High-resolution representative images from male **(S)** and female **(T)** mice showing the discrete subpopulation of CRF+ neurons co-expressing vGAT and VGLUT2 (indicated by white arrows). **(U)** Pie charts illustrate the co-expression of GABAergic markers within the total CRF+/vGLUT2+ population. In both sexes, the majority of CRF+ neurons (76–80%) are purely glutamatergic. A consistent proportion (17–18%) also express GAD2. A negligible but statistically distinct subpopulation expresses vGAT in the absence of GAD2 (as shown in S, T). This phenotype exhibited a sex difference, with a higher prevalence in males (5%) compared to females (2%), demonstrating that males possess a significantly larger fraction of CRF glutamatergic neurons carrying the vesicular GABA transporter. Scale bars: 10μm. 2-way ANOVA, Student’s t-test Chi-square test; **p<0.01.

### LHb^CRF^ neurons provide both local and long-range projections

To characterize the anatomical outputs of the LHb^CRF^ population, we mapped their efferent projections using AAV5-Syn-FLEX-ChrimsonR-tdTomato in CRH-Cre::Ai6 mice of both sexes (Figures 5-6). We utilized ChrimsonR because its red-shifted excitation spectrum offers superior tissue penetration and reduced light scattering compared to traditional blue-light-activated opsins, providing a significantly more robust platform for mapping distal terminal fields^53,54^. The integration of the Ai6 reporter line allowed us to rigorously validate our targeting specificity by confirming that ChrimsonR-tdTomato expression was strictly restricted to GFP-labeled CRF+ neurons within the LHb boundaries. Utilizing high-resolution Zeiss AxioScan microscopy, this standardized approach enabled the identification of LHb^CRF^ axonal projections within the LHb itself, as well as several key long-range targets.These distal projections included the LH/LHA, ZI, SC, PAG, VTA, substantia nigra pars compacta (SNC), as well as the dorsal raphe (DR), RMTg, and median raphe (MnR). These projection patterns were observed across both male and female mice (Figures 5–6). Our findings were further validated using two additional anterograde tracing strategies in males (AAV8-DIO-mCherry) and females (AAV5-EF1a-ChR2-EYFP). These complementary methods confirmed the axonal projections identified by our ChrimsonR approach within the LHb and major midbrain structures, including the RMTg, PAG, and SNC (Supplementary Figures 8-9). Together, these data establish that LHb^CRF^ neurons form a widespread network reaching multiple diencephalic, midbrain, and hindbrain centers.

**Figure 5:**
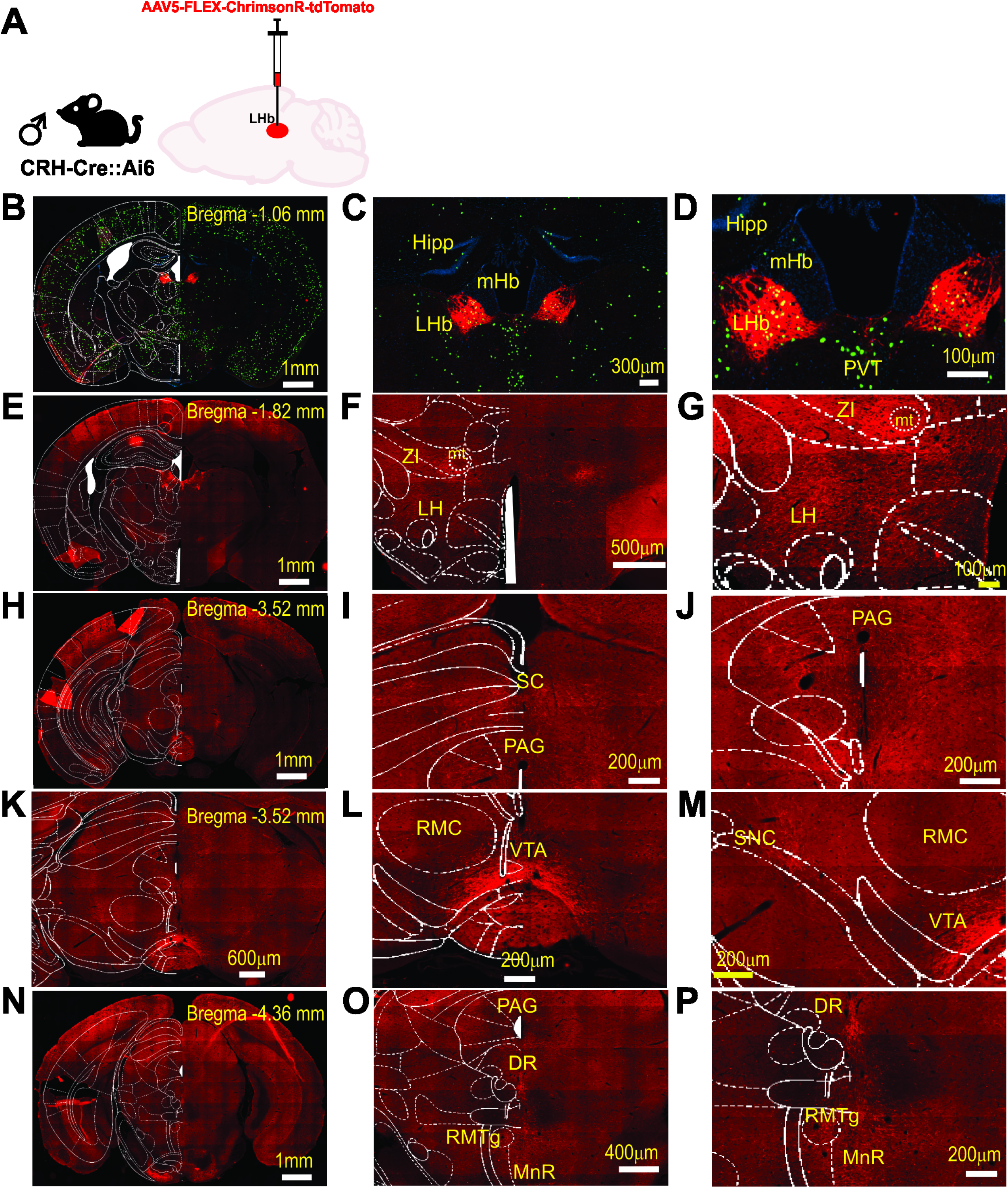
Anterograde Mapping of LHb CRF+ Neuronal Projections in male mice. **(A)** Schematic of the viral approach: a Cre-dependent anterograde vector (AAV5-syn-FLEX-ChrimsonR-tdTomato) was injected into the LHb of male CRH-Cre::Ai6 mice (n=2). (**B**–**D**) Low-magnification hemicoronal view (**B**) and high-magnification images (**C**, **D**) of the LHb at Bregma −1.06 mm. Images show the colocalization of endogenous GFP (from the Ai6 reporter line) and ChrimsonR-tdTomato, confirming the selective transduction of LHb CRF+ neurons. (**E**–**G**) Representative hemicoronal (E) and high-magnification (F, G) images illustrating tdTomato-labeled axonal processes originating from LHb CRF+ neurons within the lateral hypothalamus (LH) and zona incerta (ZI) at Bregma −1.82mm. (**H**–**J**) Midbrain Projections. Representative hemicoronal (H) and high-magnification (I, J) images showing dense tdTomato-labeled axonal innervation within the superior colliculus (**SC**) and periaqueductal gray (**PAG**) at Bregma −3.52mm. (**K**–**M**) Dopaminergic and Monoaminergic Nuclei. Representative hemicoronal (K) and high-magnification (L, M) views of labeled LHb CRF+ fibers within the ventral tegmental area (VTA) and substantia nigra pars compacta (SNc) at Bregma −3.52 mm. (**N**–**P**) Hindbrain and Raphe Nuclei. Representative hemicoronal (N) and high-magnification (O, P) images illustrating projection patterns within the periaqueductal gray (PAG), dorsal raphe (DR), rostromedial tegmental nucleus (RMTg), and median raphe (MnR) at Bregma −4.36mm. Abbreviations: PVT, paraventricular nucleus of thalamus; RMC, red nucleus, magnocellular part; mHb, medial habenula; mt, mammillotegmental tract; Hipp, hippocampus. Scale bars: B, E, H, N = 1mm; K = 600μm; F = 500μm; O = 400μm; C = 300 μm; I, J, L, M, P = 200μm; D, G= 100μm.

**Figure 6:**
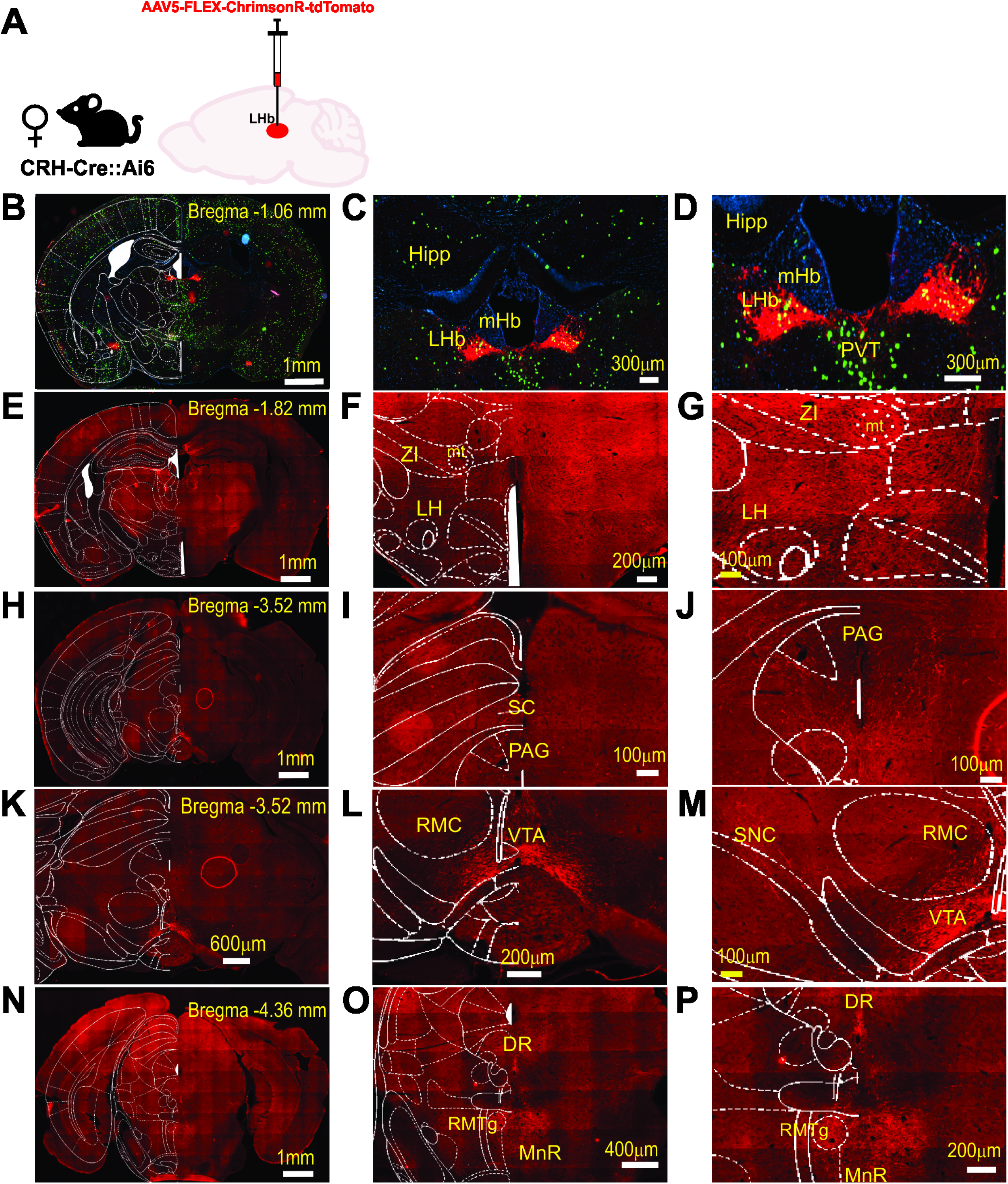
Anterograde Mapping of LHb CRF+ Neuronal Projections in female mice. **(A)** Schematic of the viral approach: a Cre-dependent anterograde vector (AAV5-syn-FLEX-ChrimsonR-tdTomato) was injected into the LHb of female CRH-Cre::Ai6 mice (n=2). (**B**–**D**) Low-magnification hemicoronal view (**B**) and high-magnification images (**C**, **D**) of the LHb at Bregma −1.06 mm. Images show the colocalization of endogenous GFP (from the Ai6 reporter line) and ChrimsonR-tdTomato, confirming the selective transduction of LHb CRF+ neurons. (**E**–**G**) Representative hemicoronal (E) and high-magnification (F, G) images illustrating tdTomato-labeled axonal processes originating from LHb CRF+ neurons within the lateral hypothalamus (LH) and zona incerta (ZI) at Bregma −1.82mm. (**H**–**J**) Midbrain Projections. Representative hemicoronal (H) and high-magnification (I, J) images showing dense tdTomato-labeled axonal innervation within the superior colliculus (**SC**) and periaqueductal gray (**PAG**) at Bregma −3.52mm. (**K**–**M**) Dopaminergic and Monoaminergic Nuclei. Representative hemicoronal (K) and high-magnification (L, M) views of labeled LHb CRF+ fibers within the ventral tegmental area (VTA) and substantia nigra pars compacta (SNc) at Bregma −3.52 mm. (**N**–**P**) Hindbrain and Raphe Nuclei. Representative hemicoronal (N) and high-magnification (O, P) images illustrating projection patterns within the periaqueductal gray (PAG), dorsal raphe (DR), rostromedial tegmental nucleus (RMTg), and median raphe (MnR) at Bregma −4.36mm. Abbreviations: PVT, paraventricular nucleus of thalamus; RMC, red nucleus, magnocellular part; mHb, medial habenula; mt, mammillotegmental tract; Hipp, hippocampus. Scale bars: B, E, H, N = 1mm; K = 600μm; O = 400μm; C, D = 300 μm; F, L, P = 200μm; G, I, J, M =100μm.

### Behavioral correlates of LHb^CRF^ neuronal activity

To establish causal links between LHb^CRF^ neuronal activity and behavior, we employed a Cre-dependent chemogenetic approach using excitatory Designer Receptors Exclusively Activated by Designer Drugs (DREADDs). Mice across multiple cohorts underwent behavioral testing (7-day intervals) in one of two sequences: light-dark test (LDT) followed by elevated zero maze (EZM), or open field test (OFT) followed by conditioned place preference (CPP) and then visual looming shadow test (VLST). The inclusion of EZM, LDT, and OFT was crucial for interpreting CPP and VLST data by controlling for potential non-specific effects of Gq-DREADD activation of LHb^CRF^ neurons on locomotion and general anxiety. To selectively activate LHb neurons expressing CRF, we employed a Cre-dependent chemogenetic approach using two different constructs of DREADDs. In adult male and female CRH-Cre mice (postnatal day 42 [P42]), we performed bilateral stereotaxic injections into the LHb of an adeno-associated viral (AAV) vector encoding an excitatory Gq-coupled DREADD. Specifically, we utilized AAV8-hSyn-DIO-hM3D(Gq)-mCherry in CRH-Cre mice to drive Cre-dependent expression of hM3D(Gq) tagged with mCherry (Supplementary Figure 10).

Given the expression of the tdTomato reporter in CRH-Cre::Ai14 mice, we used an alternative Cre-dependent viral construct, AAV8-hSyn-DIO-HA-hM3D(Gq)-IRES-mCitrine, to avoid spectral overlap and ensure clear visualization of DREADD-expressing cells (Figure 7). To activate LHb^CRF^ neurons expressing Gq-DREADDs in CRH-Cre or CRH-Cre::Ai14 mice, mice received i.p. injections of 0.3mg/kg JHU37160 or vehicle (saline) 30 min prior to behavioral assay, at least four weeks post-surgery (consistent with our previous study^55^). During CPP conditioning, mice received JHU37160 or saline prior to the second of two daily conditioning sessions (days 2-3), but not during baseline evaluation (day 1) or testing (day 4). We confirmed that Gq-DREADD expression was restricted to the LHb^CRF^ neurons by mCherry expression (Supplementary Figure 10B) or HA-hM3Dq immunostaining (Figure 7C-E) and that bath application of JHU37160 (100nM) significantly increased neuronal excitability of LHb^CRF^ neuron expressing DREADD construct (Figure 7B).

**Figure 7:**
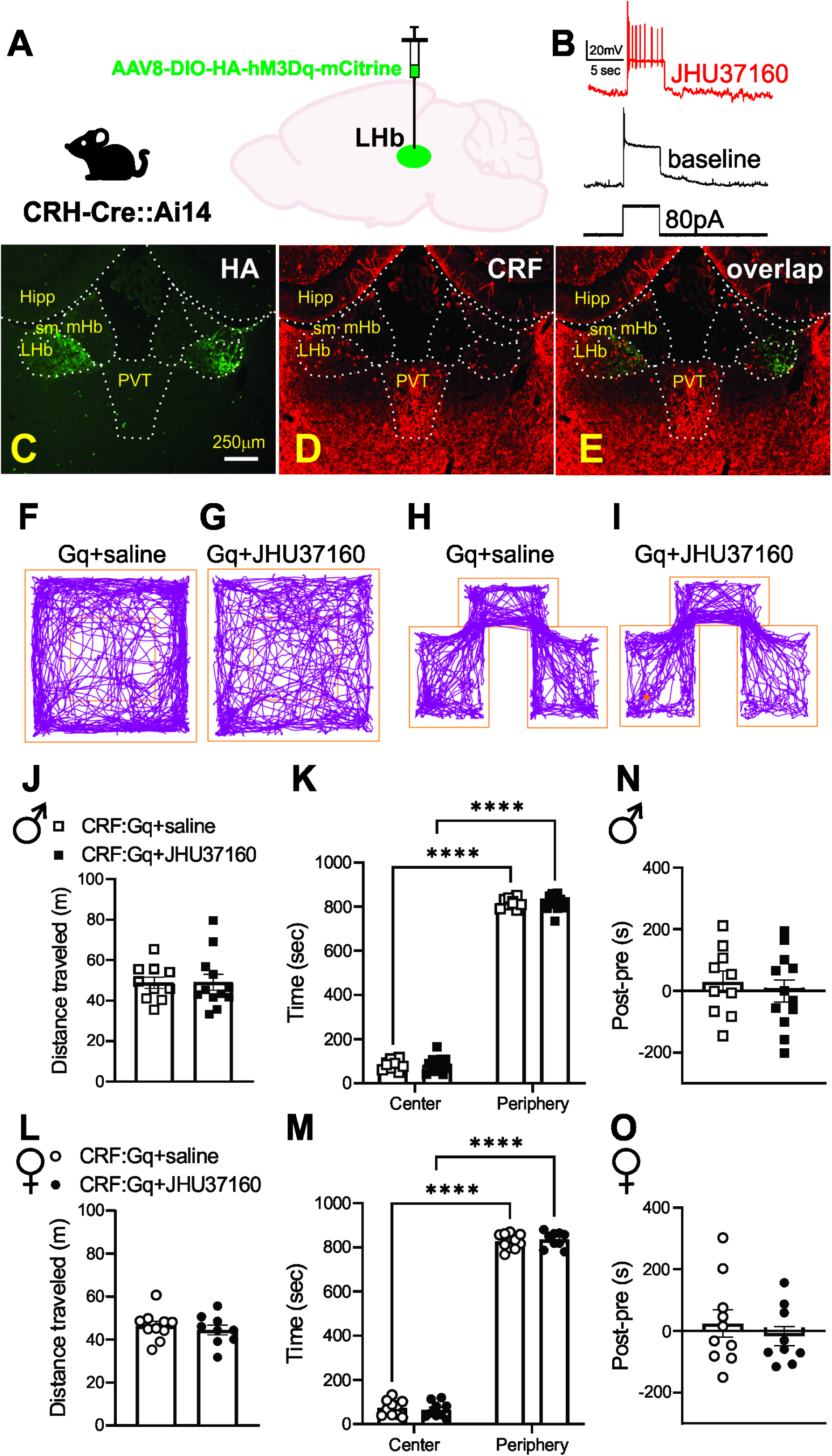
Chemogenetic activation of LHb^CRF^ neurons failed to elicit changes in locomotor activity or anxiety-like behavior in the open field test (OFT) and did not induce place preference or aversion in the conditioned place preference Test (CPP). **A** Injection schematic: AAV8-DIO-HA-hM3Dq-mCherry into the LHb of a CRH-Cre::Ai14 mouse. **B** Representative traces of action potential generation in response to an 80pA current step in an LHb^CRF^ neuron expressing hM3Dq+ before and after bath application of 100 nM JHU37160. **C-E** Fluorescent images of a coronal LHb section from a CRH-Cre::Ai14 mouse injected with AAV-DIO-HA-hM3Dq, stained with anti-HA antibody, confirming co-expression of tdTomato (red) and HA-hM3Dq (green) in LHb^CRF^ neurons, scale bar 250μm. **F-I** Representative track plots showing mouse movement in the OFT (**F-G**) and in the saline-/JHU37160-paired and unpaired compartments of the CPP (**H-I**) for saline- or JHU37160-injected CRH-Cre::Ai14 Gq-DREADD mice. **J-O** show behavioral data in OFT and CPP in male (squares) and female (circles) mice treated with either saline (open symbols) or JHU37160 (filled symbols). **J-K** and **L-M** Quantification of OFT performance revealed no significant differences between JHU37160- and saline-treated mice of either sex in total distance traveled or time spent in the center or periphery of the arena. **N-O** CPP analysis showed no evidence of place preference in JHU37160-paired CRH-Cre::Ai14 Gq-DREADD mice, comparable to saline-paired ones. Abbreviations: *LHb*, lateral habenula; *mHb*, medial habenula; *Hipp*, hippocampus; *sm*, stria medullaris; *PVT*, paraventricular nucleus of the thalamus. Scale bar: C = 250 μm. *****p<0.0001, two-way ANOVA.

### Chemogenetic activation of LHb^CRF^ neurons did not induce anxiety-like or risk-taking behaviors

In EZM and LDT, mice generally prefer to spend more time in the closed arms of the EZM or the dark chamber of the LDT box. However, they may also be motivated to explore the open arms of the EZM or the bright area of the LDT apparatus. Increased exploration of the open arms and light chamber is believed to reflect lower anxiety/more novelty seeking/risk-taking behaviors in mice. As shown in example traces of automated tracking of mouse movement in the EZM and LDT in Supplementary Figure 10C-D, activation of LHb^CRF^ neurons by JHU37160 in CRH-Cre mice expressing Gq-DREADD did not alter time spent in the open arms of EZM or the light chamber of LDT [Supplementary Figure 10E-G, EZM: Male (n=5-7/group); arm effect: F (1, 20) = 280.1, p<0.0001, JHU37160 effect: F (1, 20) = 0.031, p=0.86, arm x JHU37160 interaction: F (1, 20) = 0.34, p=0.56; Supplementary Figure 10H-J, EZM: Female (n=5-6/group); arm effect: F (1, 18) = 30.39, p<0.0001, JHU37160 effect: F (1, 18) = 0.00039, p=0.98, arm x JHU37160 interaction: F (1, 18) = 0.35, p=0.55; Supplementary Figure 10K-M, LDT: Male (n=5-7/group); chamber effect: F (1, 20) = 98.61, p<0.0001, JHU37160 effect: F (1, 20) = 0.07734, p=0.78, chamber x JHU37160 interaction: F (1, 20) = 0.34, p=0.56; Supplementary Figure 10O-P, LDT: Female (n=5-6/group); chamber effect: F (1, 18) = 34.68, p<0.0001, JHU37160 effect: F (1, 18) = 9.026e-005, p=0.99, chamber x JHU37160 interaction: F (1, 18) = 1.266, p=0.27, two-way ANOVA].

### Chemogenetic activation of LHb^CRF^ neurons did not induce CPP

To control for Gq-DREADD effects on locomotor activity and general anxiety with EZM and LDT in CRH-Cre::Ai14 lines, we performed OFT where we consistently observed that chemogenetic excitation of LHb^CRF^ neurons by JHU37160 in CRH-Cre::Ai14 mice did not affect locomotor activity or promote anxiety-like behaviors in the OFT in either sex as shown in example traces in Figure 7F-G. CRH-Cre::Ai14 mice expressing Gq-DREADD injected with either JHU37160 or saline spent significantly less time in the center of the apparatus compared to the edges (periphery) and their total distance traveled was not altered by our chemogenetic manipulation [Figure 7J-K, OFT: Male (n=10-12/group); region effect: F (1, 40) = 6656, p<0.0001, JHU37160 effect: F (1, 40) = 0.000, p=0.99, region x JHU37160 interaction: F (1, 40) = 0.018, p=0.89; Figure 7L-M, OFT: Female (n=9-10/group); region effect: F (1, 34) = 4476, p<0.0001, JHU37160 effect: F (1, 18) = 0.000, p=0.99, region x JHU37160 interaction: F (1, 34) = 0.5729, p=0.45, two-way ANOVA]. We then performed CPP to evaluate whether chemogenetic activation of LHb^CRF^ neurons is rewarding. We found that CRH-Cre::Ai14 mice expressing Gq-DREADD injected with JHU37160 during place conditioning spent equal time in JHU37160-paired and unpaired sides as seen in CRH-Cre::Ai14 mice expressing Gq-DREADD that received only saline during conditioning (Figure 7H-I and 7N,O).

### Chemogenetic activation of LHb^CRF^ neurons increased passive action-locking behavioral responses to threat

Finally, we tested whether chemogenetic activation of LHb^CRF^ neurons biases threat-evoked responses towards active (escape) or passive (action-locking/freezing) defensive behaviors and further assessed how this activation influences the temporal dynamics of these responses, including escape latency, action-locking duration, and post-escape shelter stayin mice (Figure 8). In CRH-Cre::Ai14 mice expressing Gq-DREADDs, administration of JHU37160 (0.3 mg/kg) significantly increased the frequency and duration of passive action-locking/freezing responses in both males and females compared to saline-treated controls (Figure 8B: effect of JHU37160: p < 0.05, Chi-squared tests; 10C: JHU37160 effect: F (1, 31) = 8.027, p < 0.01, sex effect: F (1, 31) = 0.2792, p=0.601, JHU37160 x sex interaction: F (1, 31) = 0.1724, p=0.68). Interestingly, we observed that activation of LHb^CRF^ neurons significantly prolonged the latency to initiate escape in male mice but had no such effect in females (Figure 8D; JHU37160 effect: F(1, 35) = 9.030, p < 0.01; sex effect: F(1, 35) = 0.062, p = 0.80; JHU37160 x sex interaction: F(1, 35) = 3.805, p = 0.05). Further analysis revealed sex-specific differences in post-threat behavior. Gq-activation in females resulted in a significantly prolonged stay within the shelter following escape (Figure 8E; JHU37160 effect: F (1, 34) = 1.707, p=0.2001; sex effect: F (1, 34) = 6.192, p<0.05; JHU37160 x sex interaction: F (1, 34) = 7.991, p<0.01). Collectively, these findings demonstrate that LHb^CRF^ neurons selectively increase the prevalence and duration of passive defensive strategies in both sexes, with sex-specific effects manifesting as differences in escape latency and shelter-seeking duration.

**Figure 8:**
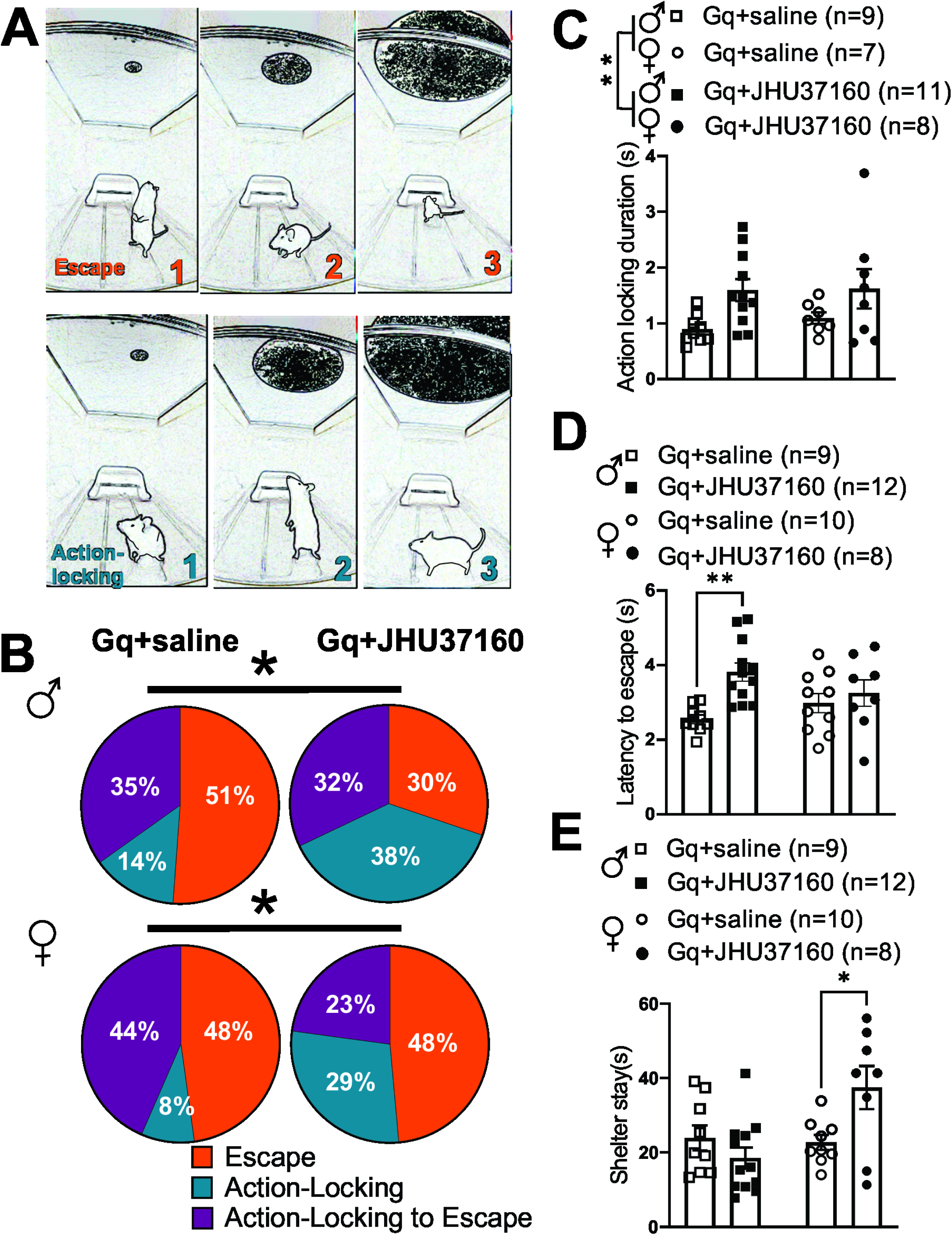
Chemogenetic activation of LHb^CRF^ neurons shifts defensive strategies toward passive action-locking in the Visual Looming Shadow Test (VLST). **(A)** Representative image sequences illustrating the two primary defensive responses to a visual looming stimulus: Escape (top row) and action-locking (bottom row). **(B)** Pie charts illustrating the percentage of trials resulting in action-locking (teal) vs. escape (orange) as well as action-locking to escape (purple) behaviors in male and female CRH-Cre::Ai14 mice expressing hM3Dq. In both sexes, administration of the DREADD agonist JHU37160 significantly increased the prevalence of action-locking behaviors. **(C–E)** show behavioral parameters in VLST in male (squares) and female (circles) mice treated with either saline (open symbols) or JHU37160 (filled symbols). **(C)** Action-locking duration: Activation of LHb^CRF^ neurons resulted in an overall increase in the duration of action-locking in both male and female mice. Of note, three mice (one male Gq+JHU, one female Gq+Saline, and one female Gq+JHU) did not exhibit any action-locking behavior across all trials. **(D)** Latency to escape: Activation of LHb^CRF^ neurons significantly increased latencies to escape in males. **(E)** Shelter Stay Duration: Post-escape behavior analysis revealed that chemogenetic activation of LHb^CRF^ neurons significantly prolonged the duration spent within the shelter specifically in female mice. Data are presented as mean ±SEM. # p=0.08, *p<0.05, **p<0.01, Chi-square test, 2-way ANOVA.

### LHb^CRF^ neurons exhibit higher baseline spontaneous activity in male mice compared to female mice

To further investigate baseline physiological differences in **LHb^CRF^**activity, we further evaluated basal levels of spontaneous activity and neuronal excitability of LHb^CRF^ neurons (labeled with tdTomato) in CRH-Cre::Ai14 male and female mice (Figure 9A). We observed that male LHb^CRF^ neurons exhibited higher levels of spontaneous activity with more LHb^CRF^ neurons in tonic and bursting mode of activity than female LHb^CRF^ neurons (Figure 9B). In contrast, LHb^CRF^ neuronal excitability in response to depolarization did not differ between male and female CRH-Cre::Ai14 mice (Figure 9C) (Figure 9B, male: n=49 cells/22 mice, female: n=41 cells/9 mice in pie charts, *p<0.05, Chi squared tests; Figure 9C, male: n=42 cells/22 mice, female: n=30 cells/9 mice in excitability, effect of sex: F (1, 740) = 0.096, p=0.75, 2-way ANOVA).

**Figure 9:**
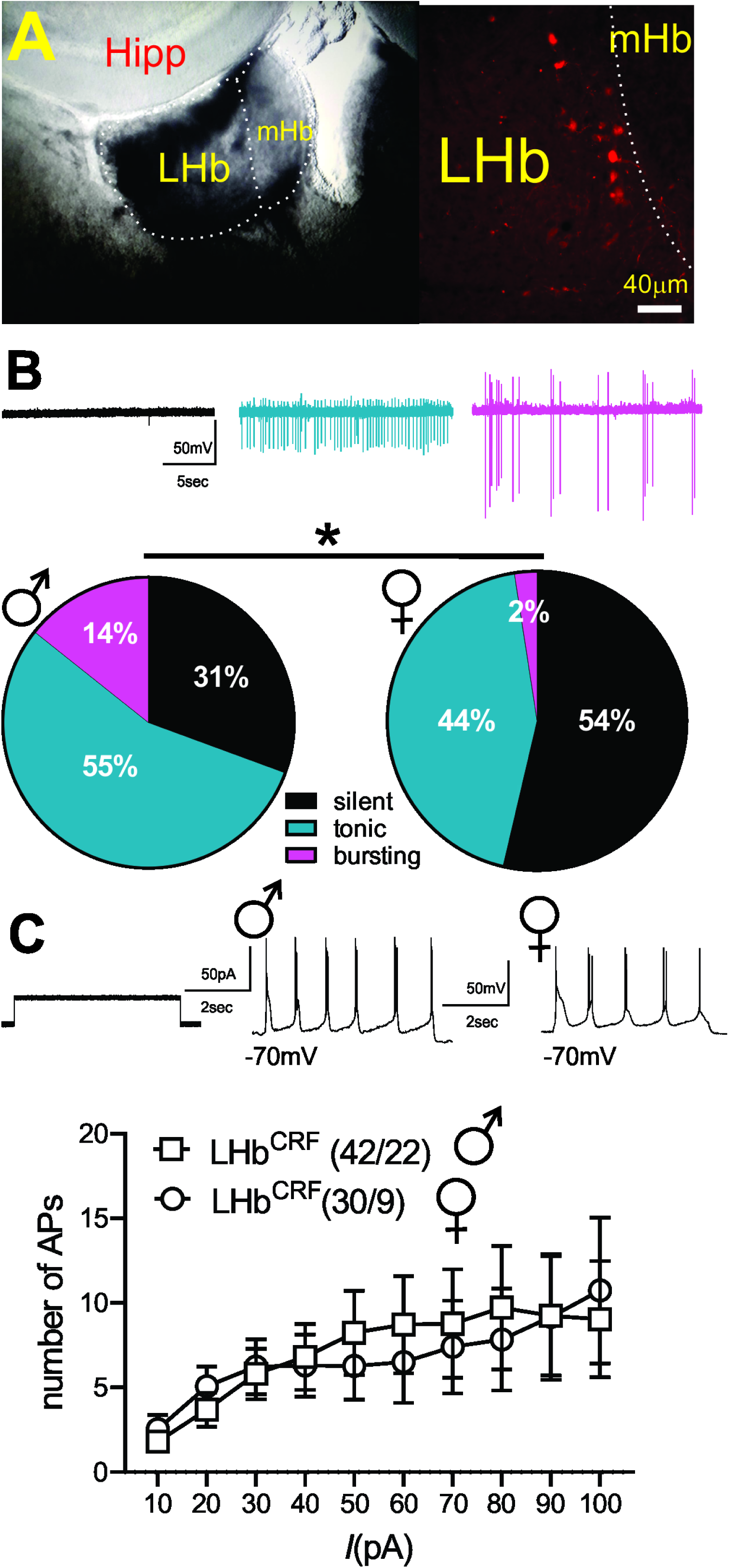
LHb^CRF^ neurons in male CRH-Cre::TdTomato mice exhibit greater spontaneous tonic and bursting activity than in females. **A** Coronal habenula slice images from a CRH-Cre::tdTomato mouse, visualized using IR-DIC (left) and fluorescence (right; scale bar = 40 μm). **B** Representative traces and pie charts illustrating spontaneous neuronal activity and the percentage of silent (black), tonic (blue), or bursting (pink) activity patterns in tdTomato-labeled LHb^CRF^ neurons from male (n=49 cells/22 mice) and female (n=41 cells/9 mice) CRH-Cre::TdTomato mice (voltage-clamp, cell-attached recordings at V=0 mV). **C** Action potential (AP) generation in response to depolarizing current steps in tdTomato-labeled LHb^CRF^ neurons, recorded under conditions of intact synaptic transmission, with representative traces from male (black open squares, n=42 cells/22 mice) and female (black open circles, 30 cells/9 mice) CRH-Cre::TdTomato mice. *p<0.05; Chi-square test. Scale bars: A = 40 μm.

### LHb^CRF^ neurons in females form stronger local excitatory connections within the LHb

To functionally validate our anterograde tracing and confirm the existence of a local microcircuit, we employed an optogenetic approach in CRH-Cre::Ai6 mice. By injecting AAV5-Syn-FLEX-ChrimsonR-tdTomato into the LHb, we specifically targeted and excited LHb^CRF^ neurons, identifying them via endogenous GFP labeling (Figure 10A–B). We performed whole-cell patch-clamp recordings from neighboring GFP-negative (non-CRF) neurons while optically stimulating ChrimsonR-expressing CRF+ cells and their local terminals. These experiments revealed robust light-evoked excitatory postsynaptic currents (oEPSCs), providing functional evidence of a direct intra-habenular glutamatergic microcircuit. We verified that these connections are monosynaptic, as the application of TTX abolished most of the oEPSCs, while subsequent 4-AP treatment recovered the currents nearly to baseline levels (Figure 10C). Recordings at a holding potential of +10 mV (the reversal potential for AMPA and NMDA receptor-mediated currents) showed no detectable oIPSCs or oEPSCs. However, at +40 mV, mixed oEPSCs were reliably evoked in both male and female mice (Figure 10D–E). Our optogenetic dissection further revealed that this local circuitry is sexually dimorphic.

**Figure 10:**
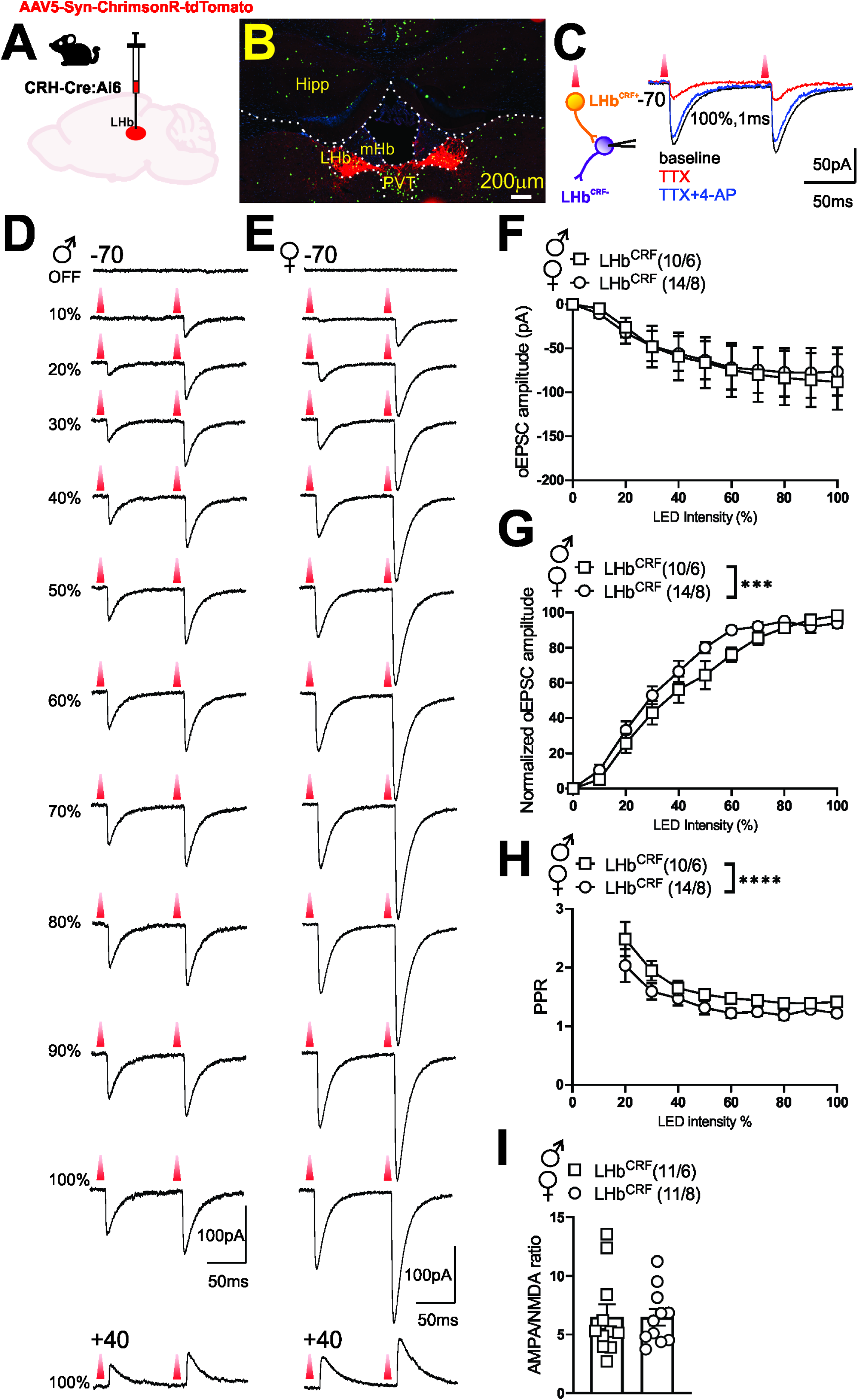
LHb CRF+ Neurons form local monosynaptic glutamatergic connections with non-CRF neurons. **(A)** Schematic illustrating the injection of a Cre-dependent anterograde vector (AAV5-Syn-FLEX-ChrimsonR-tdTomato) into the LHb of CRH-Cre::Ai6 mice to dissect local synaptic connectivity. **(B)** High-magnification image demonstrating the restricted expression of ChrimsonR-tdTomato within LHb CRF+ neurons and their local terminal fields. **(C)** Schematic and representative traces of optically evoked excitatory postsynaptic currents (oEPSCs) recorded from a non-CRF (GFP-negative) LHb neuron. Inward currents recorded at −70mV (black trace) were almost completely abolished by the sodium channel blocker TTX (1μM; red trace) and subsequently rescued by the addition of the potassium channel blocker 4-AP (50μM; blue trace), confirming the monosynaptic nature of the connection. **(D, E)** Representative traces of paired-pulse oEPSCs recorded at −70mV from male (D) and female (E) mice across increasing LED intensities (10–100%). Bottom traces illustrate mixed AMPA- and NMDA-mediated oEPSCs recorded at +40mV at maximal (100%) LED intensity. **(F)** Mean oEPSC amplitudes (measured from the first pulse) across increasing LED intensities for male (squares) and female (circles) mice. **(G)** Normalized oEPSC amplitudes expressed as a percentage of the minimum and maximum response per cell to account for variability in viral expression. **(H)** Paired-Pulse Ratio (PPR). Quantification of the PPR (P2/P1) across LED intensities male (squares) and female (circles) mice. **(I)** AMPA/NMDA Ratios. Mean AMPA/NMDA ratios recorded from LHb CRF^−^ neurons in male (squares) and female (circles) mice, showing no significant sex differences in postsynaptic receptor composition. In all graphs, n =cells/mice. Data are presented as mean±SEM. Abbreviations: *Hipp*, hippocampus; *mHb*, medial habenula; *PVT*, paraventricular nucleus of the thalamus. Scale bars: B = 200μm. ***p<0.001, ****p<0.0001; Student’s t-test and 2-way ANOVA.

While the raw input/output (I/O) curves across varying LED intensities showed no significant difference between sexes (Figure 10F; sex effect: F(1, 242) = 0.071, p = 0.789), normalized data, which controlled for variability in viral expression by scaling responses between minimum and maximum synaptic values, revealed a significant leftward shift in females (Figure 10G; sex effect: F(1, 242) = 11.27, p < 0.001). This indicates that female LHb^CRF^ neurons form significantly stronger local excitatory connections than their male counterparts. This enhanced synaptic gain in females is driven by a higher probability of presynaptic glutamate release, as evidenced by a significantly lower paired-pulse ratio (PPR) compared to males (Figure 10H; sex effect: F(1, 190) = 16.23, p < 0.0001). In contrast, we found no sex differences in the AMPA/NMDA ratio (Figure 10I; p = 0.975), suggesting that this sexual dimorphism is mediated by presynaptic release dynamics rather than differences in postsynaptic receptor composition or plasticity. Collectively, these data establish a high-gain local excitatory drive in females that may underlie their distinct behavioral responses to threat.

## Discussion

The LHb’s broad involvement in various adaptive behaviors and neuropsychiatric disorders, including depression, schizophrenia, Parkinson’s disease, and attention-deficit hyperactivity disorder, continues to fuel significant clinical interest^56–64^. Despite displaying relatively uniform electrophysiological properties, LHb neurons exhibit substantial heterogeneity in their transcriptional profiles, afferents, and projections^6,7,65,66^. Furthermore, neuromodulation profoundly shapes LHb neuronal activity through mechanisms such as the synaptic co-release of multiple neuromodulators alongside neurotransmitters, the expression of diverse neuromodulatory receptors, and the integration of various neuromodulatory signals, potentially contributing to the functional specializations of distinct LHb neuron subpopulations^2^. Given this less explored facet of LHb function and building on our previous demonstration of the LHb’s responsiveness to the key stress neuromodulator CRF^35,36,38^, we specifically investigated the origin of these previously uncharacterized CRF inputs to the LHb. Our investigation revealed several brain regions including a distinct subpopulation of LHb neurons that express CRF (LHb^CRF^). LHb^CRF^ neurons universally co-express VGLUT2. Within this population, we identified a notable subset of VGLUT2+/GAD2+ neurons, which together could serve as a source of both local intra-habenular microcircuits and widespread distal neuromodulatory inputs to downstream targets. Using gain-of-function chemogenetic strategies, we further demonstrate that LHb^CRF^ neurons may play a pivotal role in biasing behavioral selection toward passive defensive responses (i.e., action-locking behaviors in VLST). While this effect is observed across both sexes, our findings suggest that sexually dimorphic cellular and synaptic mechanisms may underlie these shared passive behavioral outputs, while also driving distinct sex differences in the temporal dynamics of the response.

Retrograde mapping in adult CRH-Cre mice identified the PRT, PAG, PVT, and the LHb itself as the primary sources of CRF inputs to the LHb. These major drivers are complemented by significant contributions from the MRN and SC, as well as consistent inputs from the PVH, PH, LH/LHA, BNST, and LPO. While many of these pathways including local intra-habenular microcircuits and long-range limbic inputs are well-documented^1,2,9,11,67–70^, the recently identified PVT-LHb connection specifically highlights the evolving understanding of thalamic influence on this region. Notably, we found no existing literature documenting a direct connection between the PRT and the LHb, suggesting this may be a previously uncharacterized pathway^69^. We acknowledge that our retrograde strategy likely provides a conservative or “limited view” of the total CRF inputs, as certain pathways may be less amenable to retrograde transduction via rgAAV vectors. The lack of prior anatomical evidence for the PRT-LHb connection underscores the need for further investigation to validate this finding and explore the functional role of these PRT^CRF^ neurons in LHb-mediated behaviors.

We then specifically focused on the newly discovered subpopulation of CRF-expressing LHb neurons (LHb^CRF^), mapping their axonal projection targets and characterizing their physiological and behavioral profiles in both sexes. Our RNAscope analysis revealed that these neurons are primarily concentrated in the rostral LHb, exhibiting a distinct spatial organization: a dense cluster resides in the medial LHb (LHbM) bordering the medial habenula, while a significant subpopulation extends into the lateral LHb (LHbL). While this topographical distribution is identical between sexes, we found that males possess a significantly higher absolute number of LHb^CRF^ neurons than females.

Transcriptional profiling confirmed that 100% of these neurons express VGLUT2. Approximately 17-18% co-express GAD2, while only a negligible minority (primarily in males) co-express vGAT. Crucially, our optogenetic data provides functional evidence for this excitatory topography; light-evoked currents recorded from local non-CRF neurons confirmed that these rostral LHb^CRF^ neurons form a functional intra-habenular circuit, with females exhibiting significantly stronger synaptic connections. Notably, although a GAD2+/VGLUT2+/vGAT- population has been reported to provide local inhibition within the LHb^8^, our functional experiments demonstrated that stimulation of LHb^CRF^ neurons evoked purely glutamatergic oEPSCs in local targets with no detectable inhibitory component. This suggests that the GAD2-expressing LHb^CRF^ subset likely corresponds to the distinct excitatory class recently defined by Quina et al. (2020), which serves as a glutamatergic output pathway to the mesopontine tegmentum^71^. Therefore, we conclude that the LHb^CRF^ population is functionally excitatory, and the GAD2-positive fraction likely contributes to long-range excitatory projections rather than local inhibition. While a minor fraction of VGLUT2+ LHb^CRF^ neurons were found to be vGAT+/GAD2-, potentially providing an inhibitory substrate (particularly in males), we did not encounter these functional connections in our recordings, as no inhibitory currents were observed. Regarding our synaptic validation, it is of note that TTX did not completely abolish oEPSCs. While this treatment typically blocks all mono- and polysynaptic currents, the use of ChrimsonR can result in sufficient terminal depolarization to trigger neurotransmitter release even in the absence of action potential, leaving a small residual oEPSC. Nevertheless, the recovery of the majority of the oEPSC amplitude following 4-AP treatment confirms the monosynaptic nature of this excitatory circuit.

Complementing this local microcircuitry, our anterograde tracing studies identified widespread LHb^CRF^ axonal projections not only within the LHb but also to several key long-range targets. These distal projections include hypothalamic/thalamic structures including LH/LHA and ZI, midbrain structures including SC, PAG, VTA and SNC and hindbrain including DR, RMTg, and MnR. Notably, all of these regions have been previously reported as primary output targets of LHb neurons in male and female mice^2,69,70,72^. Collectively, this diverse projection profile, paired with a potent local excitatory microcircuit, positions LHb^CRF^ neurons as a central hub capable of orchestrating complex behavioral and physiological responses through widespread influence on midbrain and hindbrain nuclei.

The expression of VGLUT2 in LHb^CRF^ neurons, coupled with the lack of parvalbumin expression, could indicate that LHb^CRF^ neurons are a distinct subpopulation separate from LHb^PV^ neurons. This finding also excludes the possibility that LHb^CRF^ neurons are a component of the LHb^PV^ GABAergic interneurons responsible for local inhibition of glutamatergic neurons^11^. Our future research will focus on determining whether the intrinsic and extrinsic axonal terminals of LHb^CRF^ neurons co-release glutamate and CRF to modulate synaptic transmission within the LHb and its distal downstream targets. Furthermore, we aim to elucidate the specific behavioral significance of these dual-transmitter circuits in the context of stress and defensive strategies.

Our chemogenetic investigation reveals a role for LHb^CRF^ neurons in biasing behavioral selection toward passive strategies, specifically increasing the frequency and duration of action-locking/freezing in response to looming threats in both sexes. However, the temporal execution of these defenses is strikingly dimorphic: Gq-activation prolonged escape latencies in males, whereas in females, it resulted in a protracted post-escape shelter stay. This increased shelter occupancy in females likely reflects heightened fear levels or a persistent perception of threat, necessitating a longer period to re-establish a sense of safety before re-emerging^43^.

Our data also suggests that these distinct behavioral phenotypes may be driven by divergent physiological strategies. While males possess a significantly higher absolute number of LHb^CRF^ neurons and higher basal spontaneous activity characterized by increased tonic and bursting firing modes, their local synaptic connections are less efficient than those in females. In contrast, female LHb^CRF^ neurons exhibit lower intrinsic excitability but form more potent local excitatory connections, characterized by a higher probability of glutamate release (lower PPR).

This enhanced local synaptic gain in females likely provide the physiological substrate for their prolonged action-locking and persistent shelter-seeking behavior. We propose that the lower intrinsic excitability observed in females may represent a homeostatic compensatory response to this high-gain synaptic drive, maintaining circuit stability.

Conversely, the behavioral effects in males may rely on a higher population-level output and a higher baseline firing rate. Intriguingly, while basal spontaneous activity differed between sexes, we found no sex-specific differences in overall neuronal excitability in response to direct depolarization. This discrepancy may be explained by the high proportion of bursting neurons in males. Because bursting neurons typically elicit fewer action potentials per depolarization compared to tonically active neurons during excitability protocols, the prevalence of this firing mode in males likely offsets their higher tonic activity, resulting in similar aggregate excitability levels across sexes. While we have yet to dissect the specific synaptic weights of LHb^CRF^ projections to distal targets (e.g., PAG or RMTg), it is highly probable that these sex-dependent differences in population size, baseline firing, and local synaptic gain collectively dictate the distinct behavioral outcomes observed under threat.

The hypothalamic PVH and specific extrahypothalamic CRF neurons, including the BNST (which we found both to provide weak CRF projections to the LHb), are implicated in threat-driven defensive behaviors^73^. The looming shadow test also underscores the critical involvement of both LHb neurons^44^ and PVN^CRF^ neurons^43^ in threat-provoked defensive responses. This test exhibits strong face validity as the active fleeing (flight) or passive staying (action-locking/freezing) behaviors observed closely resemble human reactions to imminent danger. Importantly, dysregulation of these defensive mechanisms, such as rigid and persistent freezing or escape, and prolonged threat reactivity, are also key features of stress-related psychopathologies like anxiety and PTSD^74–79^.

Fiber photometric recordings of PVN^CRF^ neurons in the looming shadow test demonstrated that augmented PVN^CRF^ neuronal activity precedes escape responses elicited by a looming or advancing threat^43^. Moreover, optogenetic inhibition of PVN^CRF^ neurons attenuates escape behavior^43^. Consequently, these findings suggest that PVN^CRF^ activity encodes an anticipatory or preparatory signal for the initiation of escape. Furthermore, distinct subpopulations of LHb neurons are involved in either predicting or reflecting the selection and execution of threat-driven action-locking or escape behaviors in the looming shadow test^44^. Fiber photometry of LHb population activity revealed an increase in calcium signal time-locked with escape and a decrease time-locked with action-locking. However, miniscope calcium imaging with single-cell resolution uncovered distinct LHb neuronal clusters: some increase their activity before and during escape while others decrease their activity before action-locking, while others increase activity before action-locking while decreasing activity before escape. The former outcome is reminiscent of PVN^CRF^ neuron activity during this test. Our results suggest the possibility that LHb^CRF^ neurons represent a functionally distinct LHb subpopulation that may encode action-locking defensive behaviors in response to an imminent threat. Future miniscope calcium imaging studies can further elucidate the role of LHb^CRF^ neurons and other LHb-projecting CRF neurons in the selection and execution of appropriate defensive responses. Our previous work also demonstrated that mild traumatic brain injury (mTBI) in male and female mice shifts threat responses towards action-locking in the looming shadow test and prolongs escape latency in female mice specifically^36^. Considering that mTBI-induced LHb hyperactivity is partly driven by an increase in CRF-CRFR1 signaling within the LHb in both sexes, it is crucial to investigate how LHb^CRF^ circuits contribute to the altered defensive behaviors observed after mTBI. Overall, our findings highlight that LHb^CRF^ neuronal activation shifts mouse defensive behaviors towards action-locking in both sexes. However, this activation produces distinct temporal profiles: it delays defensive reactions (increased escape latency) in males and heightens fear responses (prolonged shelter stay) in females. These findings suggest that while the core behavioral output, passive action-locking, is shared, the underlying temporal execution and post-threat recovery may be governed by sex-specific LHb^CRF^ circuit dynamics.

Curiously, our chemogenetic activation of LHb^CRF^ neurons did not alter place preference in either sex while activation of distinct subpopulations of CRF neurons could exert either place preference (e.g., NAc^CRF^ or CeA^CRF^ neurons) or aversion (e.g., BNST^CRF^)^80^. One limitation of chemogenetic activation with sustained activation of LHb^CRF^ neurons is its lack of the millisecond-scale temporal precision offered by optogenetics. This limitation hinders a clear understanding of the immediate behavioral response to LHb^CRF^ neuronal stimulation. Therefore, future studies using behavioral assays like the real-time place preference paradigm could better address whether LHb^CRF^ neuronal stimulation evokes aversive (avoidance) or appetitive (approach) motivation.

In summary, our work has unveiled a previously uncharacterized subpopulation of LHb^CRF^ neurons that form both local microcircuits and long-range projections to the midbrain and hindbrain. Our findings establish the LHb^CRF^ population as a novel, sexually dimorphic component of the LHb’s heterogeneous circuit architecture that could bias defensive strategies toward action-locking under threat and may underlie the sex-dependent variations in escape latency and post-escape shelter seeking observed during the VLST. Further characterizing the precise functional connectivity and behavioral contributions of these neurons based on their unique projection targets represents a significant and timely direction for the field. Given that CRF neuromodulation is inherently cell-type, region, and pathway-specific, dissecting these distinct LHb^CRF^ circuits will provide essential insights into the nuanced mechanisms by which CRF orchestrates complex behavioral states and sex-specific stress responses.

## Methods

### Animals

Male and female mice, aged 9-13 weeks, were used for all experiments (viral injections at postnatal day 42 (P42); experiments commencing at 9 weeks of age). All procedures adhered to the NIH Guide for the Care and Use of Laboratory Animals and the ARRIVE guidelines 2.0 and were approved by the Uniformed Services University Institutional Animal Care and Use Committee. The following mouse strains were used: CRH-ires-CRE (B6(Cg)-*Crh^tm1(cre)Zjh^*/J, Strain #: 012704), n = 32; CRH-Cre::Ai14 [generated by crossing homozygous CRH-ires-Cre mice (n=16 breeders) with Ai14 Cre reporter mice (n=12 breeders)( B6.Cg-*Gt(ROSA)26Sor^tm^*^14^*^(CAG-tdTomato)Hze^*/J, Strain #: 007914)] or Ai6 Cre reporter mice (n=4 breeders) (B6.Cg-*Gt(ROSA)26Sor^tm6(CAG-ZsGreen1)Hze^*/J, Strain#: :007906), reporter lines (n=101), total n = 165. Mice were acquired from Jackson Laboratories at P35-P49 and allowed at least 72 hours of acclimation before experimental procedures began. Different numbers of male and female mice were used due to stock availability from the supplier and in-house breeding. For chemogenetic experiments, animals were randomly assigned to groups, and all data acquisition and analysis were performed by investigators blinded to the group assignments (Gq+saline/Gq+JHU37160). Seven mice were excluded from chemogenetic studies due to incorrect LHb targeting. Mice were group-housed in standard cages under a 12hr/12hr light-dark cycle with standard laboratory lighting conditions (lights on, 0600-1800, ∼200 lux), with ad libitum access to food and water. All procedures were performed 2-4 hours after the start of the light cycle. All efforts were made to minimize animal suffering and reduce the number of animals used throughout this study.

### Virus injections for retrograde and anterograde tracings and chemogenetics

Mice (P42) were anesthetized with 1-3% isoflurane vaporized in oxygen and secured in a stereotaxic frame. Body temperature was maintained at 37°C throughout the procedure and during recovery using a heating pad. The scalp was shaved, and a midline incision was made to expose the skull. The following viral vectors were bilaterally injected into the LHb: rgAAV-EF1a-DIO EYFP (Addgene #27056), AAV5-Syn-FLEX-rc[ChrimsonR-tdTomato (Addgene#62723), AAV5-Ef1a-DIO hChR2(E123T/T159C)-EYFP (Addgene #35509), AAV8-hSyn-DIO-mCherry (Addgene #50459), AAV8-hSyn-DIO-hM3D(Gq)-mCherry (Addgene #44361), AAV8-hSyn-DIO-HA-hM4D(Gi)-IRES-mCitrine (Addgene #50455). Viral vectors (∼50 nL/side) were infused at a rate of 1nL/sec over 5 minutes using a Nanoject III Injector (Drummond) and pulled glass pipettes into the LHb (coordinates from bregma: AP, −1.35 mm; ML, ±0.5 mm; DV, −3.0 mm). The injection pipette was left in place for 5 minutes post-infusion to allow for viral diffusion. During recovery (approximately 30-60 minutes), mice were placed in a warm cage (heating pad beneath) or under a heating lamp to prevent hypothermia. Post-operative care included continuous monitoring until purposeful movement was regained. Animal health and weight were monitored and recorded for at least 3 days post-surgery. Mice received a subcutaneous injection of 5 mg/kg carprofen (for analgesia) immediately after surgery and were monitored daily for signs of pain, with carprofen administered as needed. Viral expression was confirmed via fluorescence and/or immunohistochemistry at the conclusion of behavioral experiments. Mice lacking bilateral viral expression in the LHb were excluded from data analysis.

### In Situ Hybridization (RNAscope)

Multiplex fluorescent *in situ* hybridization (FISH) was performed using the RNAscope Fluorescent Multiplex Assay (Advanced Cell Diagnostics [ACD], Newark, CA) to detect mRNA transcripts for *Slc17a6* (VGLUT2), *Slc32a1* (VGAT), and *Gad2* (GAD2).

### Tissue Preparation

Male and female CRH-Cre::Ai14 mice were anesthetized with ketamine/xylazine (85/10 mg/kg, i.p.) and transcardially perfused with 100 ml of 1x PBS followed by 100 ml of 4% paraformaldehyde (PFA). Brains were post-fixed in 4% PFA (24 h), cryoprotected in 20% sucrose (72 h), and frozen on dry ice. Coronal sections (20μm) spanning the LHb (Bregma −0.94 to −2.30 mm; Paxinos and Franklin, 2007) were collected via cryostat (Leica CM1900) and mounted on glass slides.

### Sequential Imaging and RNAscope Assay

Because the endogenous tdTomato signal is quenched during the RNAscope protease and dehydration steps, a sequential imaging strategy was employed:

1. Pre-assay Imaging (Scan 1): Prior to the FISH assay, slides were scanned under a 20x objective (Zeiss Axioscan) to capture the endogenous tdTomato (red) signal from CRF+ neurons.
2. RNAscope Processing: Following Scan 1, coverslips were removed in 4x SSC, and sections were washed in 1x PBS. Sections were baked at 60° C, post-fixed in 4% PFA (1h at 4°C), and dehydrated in an ascending ethanol series. Following hydrogen peroxide treatment and antigen retrieval, sections were treated with Protease III.
3. Hybridization and Detection: Sections were incubated (2h at 40°C) with a multiplex probe cocktail: *Slc17a6*-C1 (Cat# 319171), *Slc32a1*-C3 (Cat# 319191), and *Gad2*-C4 (Cat# 439371). Signals were amplified using AMPs 1–3 and developed with HRP-conjugated Vivid fluorophores (1:5000 dilution): Vivid 520 (*Slc17a6*), Vivid 570 (*Slc32a1*), and Vivid 650 (*Gad2*).
4. Post-assay Imaging (Scan 2): After signal development, sections were counterstained and mounted with ProLong Gold with DAPI (Life Technologies). A second scan was performed under identical settings to capture the FISH signals.

### Image Processing and Registration

To visualize the colocalization of mRNA transcripts with CRF+ (tdTomato) neurons, Scan 1 (pre-assay) and Scan 2 (post-assay) images were aligned and merged using RNAscope HiPlex Image Registration Software v2.1 (ACD). This process allowed for the simultaneous analysis of *vGlut2*, *vGat*, and *Gad2* expression within the previously identified tdTomato-labeled CRF+ population.

### Image Analysis and RNAscope Quantification

QuPath Identification and Annotation: Quantitative analysis of CRF+ (tdTomato+) neurons and their co-expression of *Slc32a1* (vGAT), *Gad2* (GAD2), and *Slc17a6* (vGLUT2) mRNA was performed using QuPath (v0.5.0) image analysis software. The LHb was anatomically delineated in each section using the Allen Mouse Brain Atlas in conjunction with the Paxinos and Franklin Mouse Brain Atlas as reference. The LHb perimeter was manually outlined using the closed polygon annotation tool to create a defined Region of Interest (ROI) for automated analysis.

Cell Detection and Phenotypic Classification: Individual CRF+ neurons were identified and segmented within the LHb ROI using the Cell Detection tool, which utilized the tdTomato fluorescence signal for primary cell identification. Following detection, cells were systematically classified based on the presence of multiplexed RNAscope puncta. Phenotypes were categorized as:

- Single-positive: vGLUT2+, vGAT+, or GAD2+
- Dual-positive: vGLUT2+/vGAT+, vGLUT2+/GAD2+, or vGAT+/GAD2+
- Triple-positive: vGLUT2+/vGAT+/GAD2+

Anatomical Sampling and Statistical Representation: Quantification was conducted on coronal sections from CRH-Cre::Ai14 mice (n=13; 7 males, 6 females). To ensure an unbiased longitudinal assessment, cells were quantified across five distinct anterior-posterior (AP) levels of the LHb, ranging from Bregma −1.06 to −2.18mm.

Data are reported as the absolute number of CRF+ cells per section. Phenotypic distributions (e.g., percentage of dual- or triple-positive neurons) were calculated as a proportion of the total CRF+ population, noting that 100% of the identified CRF+ neurons were also positive for vGLUT2.

### Immunohistochemistry

At least three weeks following viral injection, similar to RNAScope procedure, mice were anesthetized and perfused transcardially and brain sections (20 μm) were cut using a cryostat, generating a series from Bregma 2.10 to −5.68mm AP. Sections were mounted on slides. For immunolabeling, LHb sections from CRH-Cre mice injected with rgAAV-EF1a-DIO EYFP (for parvalbumin staining) or CRH-Cre:Ai14 mice injected with AAV8-DIO-HA-hM3D(Gq)-mCitrine (for HA-tag staining) were blocked in 0.3% Triton X-100 containing 10% Normal Goat Serum (NGS) in 1x PBS for 1 hour and then incubated overnight at room temperature in carrier solution (5% NGS in 0.1% Triton X-100 in 1x PBS) with: mouse anti-parvalbumin IgG1 antibody (1:1000, Sigma, P3088) to detect parvalbumin expression in retrogradely labeled LHb^CRF^ neurons or rabbit anti-human influenza hemagglutinin (HA) antibody (1:500, Cell Signaling) to detect HA-hM3Dq in tdTomato-expressing LHb neurons.

Sections were rinsed in 1x PBS and incubated for 2 hours in Alexa Fluor® 594-labeled Goat anti-Mouse IgG1 or Alexa Fluor® 647-labeled Goat anti-Rabbit. Sections were rinsed again in 1x PBS and coverslipped with Prolong Gold mounting medium containing DAPI. Background staining was assessed by omitting the primary antibody (negative control). Brain tissue sections with known parvalbumin immunoreactivity (cortex, hippocampus) were also processed as positive control tissues. Images were captured using a Leica DMRXA Fluorescence microscope.

### Anterograde Projection Mapping and Histological Imaging

At least three weeks following viral injection, mice were anesthetized and underwent transcardial perfusion with 4% PFA, following the same standardized protocol used for RNAscope and immunohistochemistry. Brains were post-fixed, cryoprotected, and subsequently sectioned at 20 μm using a cryostat. To ensure a comprehensive mapping of projection targets, a continuous series of coronal sections was collected across the rostrocaudal axis, spanning from Bregma +2.10 to −5.68 mm. To map the efferent projection patterns of LHb^CRF^ neurons, three distinct anterograde viral strategies were employed. In our primary approach (Figures 5–6), CRH-Cre::Ai6 mice of both sexes were injected with AAV5-Syn-FLEX-ChrimsonR-tdTomato. For automated histological imaging, sections were scanned using a Zeiss Axioscan equipped with a 20x objective. This configuration permitted the simultaneous capture of the endogenous GFP (green) signal—representing the global population of CRF+ neurons via the Ai6 reporter—and the tdTomato (red) signal, marking the specific axonal projections originating from ChrimsonR-expressing CRF+ neurons in the LHb. The resulting high-resolution mosaic images were utilized to verify injection site specificity and to trace efferent fibers across multiple neuroanatomical targets along the rostrocaudal axis. Two alternative strategies were initially used to map these projections: AAV8-hSyn-DIO-mCherry injections in male CRH-Cre mice and AAV5-Ef1a-DIO-ChR2-EYFP injections in female CRH-Cre mice. While these initial experiments identified consistent projections to the LHb, RMTg, SNc, and PAG, we subsequently transitioned to the ChrimsonR-tdTomato approach in the Ai6 reporter line. This refined strategy allowed for a more rigorous, sex-balanced comparison and, when combined with the Zeiss Axioscan, provided superior sensitivity in detecting additional efferent projections. Consequently, the results from the initial ChR2 and mCherry experiments were documented as supplementary data (Supplementary Figures 8 and 9), with images captured using a Leica DMRXA fluorescence microscope.

### Quantification of Retrogradely Labeled CRF+ Neurons

To quantify the regional distribution of LHb-projecting CRF+ neurons, EYFP-labeled cell bodies were identified and mapped using the Allen Mouse Brain Atlas in conjunction with the Paxinos and Franklin Mouse Brain Atlas. Analysis was conducted on coronal sections from three CRH-Cre mice (n = 3; 2 males, 1 female). The total number of retrogradely labeled neurons analyzed per subject was 2,211 (CRH-Cre 1), 3,309 (CRH-Cre 2), and 1,750 (CRH-Cre 3). To ensure representative sampling across the rostrocaudal axis, labeled cells were counted within defined regions of interest (ROIs) across three consecutive sections (inter-section interval of 0.12 mm). Data are expressed as a percentage of the total population of labeled neurons identified across all ROIs: Percentage of Labeled Neurons = (Cells in ROI / Total Labeled Cells) X 100 Forebrain and hypothalamic ROIs included the bed nucleus of the stria terminalis (BNST; Bregma +0.14 to −0.22 mm), lateral preoptic area (LPO; Bregma +0.38 to −0.22 mm), paraventricular hypothalamic nucleus (PVH; Bregma −0.34 to −1.46 mm), and lateral hypothalamus (LHA; Bregma −0.34 to −1.34 mm). Midbrain and thalamic targets sampled included the paraventricular thalamic nucleus (PVT; Bregma −1.34 to −2.30 mm), lateral habenula (LHb; Bregma −1.06 to −2.30 mm), and posterior hypothalamus (PH; Bregma −2.06 to −2.80 mm). Posterior midbrain and brainstem structures included the pretectal area (PRT, including the medial pretectal nucleus; Bregma −2.46 to −2.80 mm), midbrain reticular nucleus (MRN; Bregma −2.54 to −3.80 mm), superior colliculus (SC; Bregma −2.80 to −3.92 mm), and the periaqueductal gray (PAG; Bregma −2.30 to − 4.04 mm). Figure 1 and supplementary Figure 1 include the data derived from this experiment.

### Ex Vivo Slice Electrophysiology and Optogenetics Slice Preparation

Electrophysiology experiments were conducted using male and female CRH-Cre::Ai14 reporter mice or CRH-Cre::Ai6 mice. For optogenetic experiments, CRH-Cre::Ai6 mice were previously injected with a Cre-dependent anterograde vector (AAV5-Syn-FLEX-ChrimsonR-tdTomato) into the LHb. To allow for robust opsin expression and trafficking to distal terminals, all physiological recordings were performed at a minimum of three weeks post-injection. Mice were deeply anesthetized with isoflurane and decapitated. Brains were rapidly dissected and placed in ice-cold artificial cerebrospinal fluid (ACSF) containing (in mM): 126 NaCl, 21.4 NaHCO3, 2.5 KCl, 1.2 NaH2PO4, 2.4 CaCl2, 1.00 MgSO4, 11.1 glucose, 0.4 ascorbic acid; saturated with 95% O2-5% CO2.

Sagittal or coronal slices (220μm) containing the LHb were cut using a vibratome (Leica; Wetzlar, Germany) and incubated in oxygenated ACSF at 34°C for at least 1 hour prior to recording. For all experiments, slices were transferred to a recording chamber and perfused with ascorbic-acid-free ACSF maintained at 28–30°C.

### General Recording Conditions

Whole-cell patch-clamp recordings were performed using an upright microscope (Olympus BX51WI) equipped with infrared-differential interference contrast (IR-DIC) microscopy. Patch pipettes (3–6 MΩ) were controlled via a MultiClamp 700B amplifier, digitized with a DigiData 1440A, and analyzed using pCLAMP 10 (Molecular Devices) and GraphPad Prism 10. Signals were filtered at 3kHz and digitized at 10kHz. Input and series resistances were monitored throughout; data were excluded if these values fluctuated by more than 10%.

### Intrinsic Excitability and Firing Patterns

To assess intrinsic properties of LHb^CRF^ neurons in male and female CRH-Cre::Ai14 mice, pipettes were filled with a potassium gluconate-based internal solution (130 mM K-gluconate, 15 mM KCl, 4 mM ATP-Na+, 0.3 mM GTP-Na+, 1 mM EGTA, and 5 mM HEPES, pH adjusted to 7.28 with KOH, osmolarity adjusted to 275-280 mOsm).

Spontaneous neuronal activity and action potential (AP) firing patterns (e.g., tonic vs. bursting) were recorded in cell-attached voltage-clamp mode (V = 0) for approximately one minute. To assess excitability, neurons were maintained at −65 to −70mV via manual DC injection. Increasingly depolarizing current steps (5s duration) were applied at +10pA intervals (ranging from +10 to +100pA) with a 20s inter-stimulus interval. The number of APs generated per step was quantified and averaged by experimental group.

### Optogenetic dissection of local LHb^CRF^ connectivity

To investigate local synaptic connections, recordings were obtained from non-CRF LHb neurons (GFP-negative) while optogenetically stimulating local CRF+ terminals within the LHb in male and female CRH-Cre::Ai6 mice (injected with AAV5-Syn-FLEX-ChrimsonR-tdTomato). For these experiments, pipettes were filled with a cesium gluconate-based internal solution (117 mM CsGlc, 2.8 mM NaCl, 5 mM MgCl_2_, 2 mM ATP-Na+, 0.3 mM GTP-Na+, 0.6 mM EGTA, and 20 mM HEPES, pH 7.28, osmolality 275–280 mOsm). Optogenetic stimulation (590nm LED; X-Cite XLED1, Lumen Dynamics) was delivered through a 40x0.8NA objective of an upright microscope (Olympus, BX51WI). Light pulses (1ms) were delivered at intensities of 0.1-1.0mW/mm^2^. To determine the input/output (I/O) relationship of the local synapses, peak AMPAR-mediated oEPSCs were recorded at a holding potential of −70mV across increasing LED intensity levels (0–100%). To control for variations in viral expression levels, data were then normalized for each neuron and expressed as a percentage of the minimum and maximal amplitude responses recorded across the intensity range.

Neurons were also held at specific potentials to further characterize synaptic inputs:

- +10mV: To confirm the absence of inhibitory currents (oIPSCs) at the glutamate reversal potential.
- +40mV: To evoke mixed AMPA- and NMDA-mediated oEPSCs.

The NMDAR-mediated component was isolated at +40mV by measuring current amplitude 20ms post-AMPAR peak (∼50ms post-LED trigger). For paired-pulse ratio (PPR) experiments, two light pulses were delivered with a 100ms inter-pulse interval and a 20s inter-trial interval. Peak amplitudes for both evoked oEPSCs (P1 and P2) were measured relative to a stable baseline, which was defined as the average current during a 20–50ms window immediately preceding the first light-evoked current. The PPR was calculated as the ratio of the peak amplitude of the second response to the first (P2/P1) at each intensity level.

### Pharmacological Validation

To confirm monosynaptic, glutamatergic transmission, a subset of experiments utilized bath application of the sodium channel blocker tetrodotoxin (TTX, 1μM), followed by the potassium channel blocker 4-aminopyridine (4-AP, 50μM) to rescue monosynaptic oEPSCs.

### Behavioral assays with chemogenetics

Four weeks post-viral injections, behavioral testing began with separate cohorts of CRH-Cre or CRH-Cre::Ai14 male and female mice expressing Gq DREADD in LHb. Mice received intraperitoneal injections of either 0.3mg/kg of the DREADD ligand, JHU37160, or vehicle (equivalent volumes of 0.9% saline) 30 min prior to behavioral testing, consistent with our previous study^55^. In the CPP test, DREADD-injected mice underwent conditioning with JHU37160 paired with either the preferred or non-preferred side of the chamber in a balanced (unbiased) manner. In these cohorts, JHU37160 or saline was administered only during conditioning days (days 2-3), not during habituation (day 1) or testing (day 4).

### Open field test (OFT)

Mice were placed in an open field arena (40 cm x 40 cm) with dim white lighting (∼5lux) and allowed to explore freely for 15 minutes. Animal movement was tracked using Any-Maze video tracking software (Stoelting Co.), recording distance traveled and time spent in the periphery and center zones.

### Elevated zero maze (EZM)

To assess anxiety and risk-taking behavior, mice were placed in an elevated zero maze and allowed to freely explore for 10 minutes with dim white lighting (∼5lux). The maze, elevated 40 cm above the floor, comprises a ring-shaped arena with four arms (5 cm x 30 cm): two enclosed arms with 20-cm high walls and two open arms. An overhead camera and Any-Maze software (Stoelting Co.) were used to track the animal’s movement, quantifying the number of entries and the time spent in both the open and closed arms, along with the total distance traveled.

### Light-Dark Test (LDT)

The light-dark box test, similar in concept to the elevated zero maze but without the height factor, consists of a 40 cm x 40 cm box divided into a dark, enclosed chamber and a bright, exposed chamber (∼750lux). While mice prefer the dark chamber, they also explore the bright area. Less time spent in the light chamber and more time in the dark is interpreted as higher anxiety. An overhead camera and Any-Maze software (Stoelting Co.) tracked the animal’s position, recording distance traveled and time spent in each chamber.

### Conditioned place preference (CPP)

We employed a three-compartment apparatus to conduct CPP testing. This apparatus featured outer compartments with distinct visual cues (e.g., circles vs. triangles walls) and a small central compartment lacking specific cues. Gates allowed mice to freely traverse the three compartments. Anymaze software was used to track the time spent in each compartment during the pre-test, conditioning, and test phases. The CPP procedure comprised: Pretest (Day 1): To assess baseline preferences, each mouse was placed in the central compartment and allowed to explore all three compartments for 20 minutes. Conditioning (Days 2-3): Mice underwent conditioning sessions wherein they were confined to one outer compartment and administered saline, followed by confinement to the opposite compartment (after a >4-hour interval) and administration of either saline or JHU37160 (30 minutes prior to confinement). The pairing of saline or JHU37160 with a specific compartment was balanced across animals. Test (Day 4): Mice were given free access to all compartments, and the time spent in the saline/JHU37160-paired compartment was measured to determine place preference. Place preference was quantified by calculating a preference score: time spent in the saline/JHU37160-paired compartment during the post-test minus time spent in that compartment during the pre-test (post-pre).

### Visual Looming Shadow test (VLST)

This test simulates a predator threat to induce defensive behaviors in mice, which manifest as either passive action-locking/freezing or active escape. The testing environment consisted of a 17 x 10 x 8.5-inch plastic arena with a 21-inch LCD monitor positioned overhead to present the visual stimulus. A 6 x 6 x 3-inch shelter was placed on one side of the arena. Behavior was recorded with a camera positioned laterally above the arena. First, mice were habituated to the arena for two days (10 minutes/day). On the test day, there was a 5-minute habituation period within the arena before stimulus presentation and then mice were presented with five looming visual stimuli, with at least a 1-minute inter-trial interval. The stimulus comprised of a 2-cm black disk expanding to 20 cm over 9 seconds, divided into three phases: 2 seconds at 2 cm, 5 seconds of expansion, and 2 seconds at 20 cm. Defensive responses were classified as: escape (active movement to the shelter), action-locking/freezing (absence of movement or repeated, discontinuous freezing) and action-locking to escape behavior (action-locking that transitions to escape in which mice exhibit an immediate freezing behavior followed by a delayed escape to the shelter). Mice were excluded as non-responders if they exhibited no discernible reaction to the looming stimulus (no freezing, no behavioral change, and failure to reach the shelter within 8 seconds). Video recordings were used to blindly score the type of defensive behavior and the latency to escape. For each subject, escape latencies, shelter stay, and action locking duration were averaged across the five looming trials; these mean values were then used for all subsequent statistical analyses and are presented as individual data points.

### Statistics and reproducibility

Data are presented as mean ± SEM. Statistical significance was assessed using unpaired two-tailed Student’s t-tests, Chi-square tests, and two-way ANOVA with Bonferroni post hoc tests. A significance threshold of P < 0.05 was used for all analyses. Statistical analyses were performed using GraphPad Prism 10.

## Data availability

Data supporting the conclusions of this study are available from the corresponding author upon reasonable request. Supplementary data associated with the figures are provided in the Supplementary Information files.

## Acknowledgements

The opinions and assertions contained herein are the private opinions of the authors and are not to be construed as official or reflecting the views of the Uniformed Services University of the Health Sciences or the Department of Defense or the Government of the United States. This work was supported by the National Institute of Mental Health (NIH/NIMH) Grant# R21 MH132136 to FSN. The funding agency did not contribute to writing this article or deciding to submit it. We gratefully acknowledge the Preclinical Behavior and Modeling Core for providing access to and support with their behavioral equipment, which was essential for the successful completion of this study.

## Author contributions

FSN and WJF designed the study. WJF, SCS, and EHT performed electrophysiology. TJW, SCG, and EHT were responsible for breeding CRH-Cre::Ai14 and CRH-Cre::Ai6 mice. SCG performed microscopy, RNAscope and immunohistochemistry, with assistance from LI and EP for RNAscope optimization.

SCG, SCS, and WJF conducted viral stereotaxic injections. WJF, EHT, MCT, and SCS carried out chemogenetic behavioral assays. CMD assisted in analysis and interpretation of behavioral data. Data analysis and figure preparation were performed by WJF, SCG, SCS, EHT, OLR, and FSN. The initial manuscript draft was written by WJF, SCG, EHT, RCA, CMD, EP, BMC, and FSN. All authors critically reviewed and approved the final version of the manuscript.

