## Supplemental figures 1-10 for "Neuroanatomical and behavioral characterization of corticotropin releasing factor-expressing lateral Habenula neurons in mice"

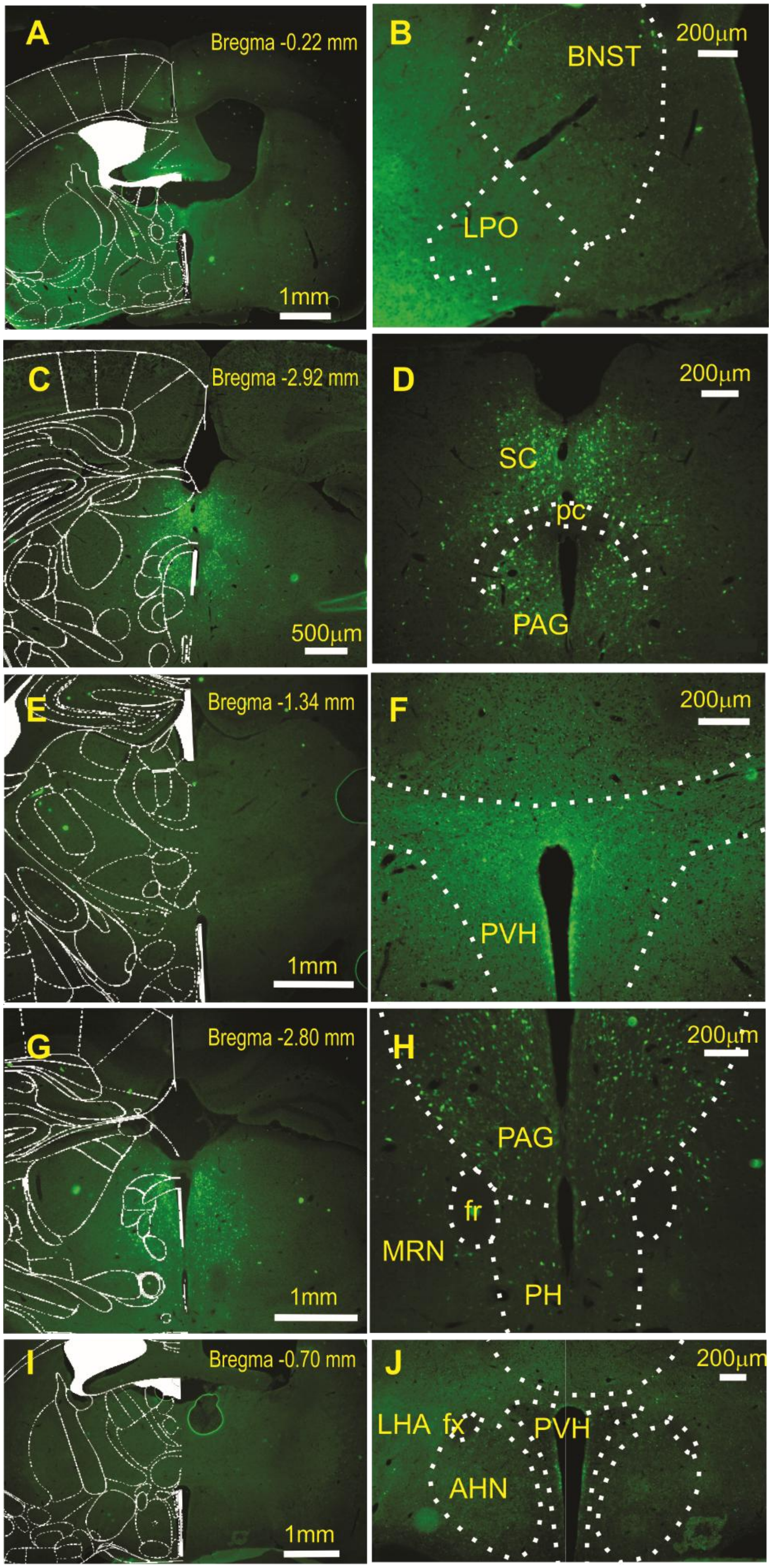

A
Bregma -0.22 mm
1mm
B
200μm
BNST
LPO
C
Bregma -2.92 mm
500μm
D
200μm
SC
pc
PAG
E
Bregma -1.34 mm
1mm
F
200μm
PVH
G
Bregma -2.80 mm
1mm
H
200μm
PAG
fr
MRN
PH
I
Bregma -0.70 mm
1mm
J
200μm
LHA
fx
PVH
AHN

**Supplementary Figure 1: Retrograde identification of CRF-expressing projections to the LHb**

**A–J)** Regional Distribution of LHb-Projecting CRF+ Neurons. Representative coronal sections showing the spatial distribution of retrogradely labeled EYFP+ neurons across the rostrocaudal axis.

- **(A, C, E, G, I)**: Low-magnification overviews integrated with mouse atlas overlays at the indicated Bregma levels (-0.22, -0.70, -1.34, -2.80 and -2.92mm) to visualize regional landmarking.
- **(B)**: High-magnification image of labeling in the bed nucleus of the stria terminalis (BNST) and lateral preoptic area (LPO).
- **(D)**: High-magnification image showing labeled neurons in the superior colliculus (SC) and periaqueductal gray (PAG).
- **(F)**: High-magnification view of the paraventricular hypothalamic nucleus (PVH).
- **(H)**: Labeled CRF+ neurons identified in the posterior hypothalamus (PH), midbrain reticular nucleus (MRN), and PAG.
- **(J)**: Distribution of labeling within the paraventricular hypothalamic nucleus (PVH) and lateral hypothalamus (LHA) near the fornix (fx).

Abbreviations: *AHN*, anterior hypothalamic nucleus; *BNST*, bed nucleus of the stria terminalis; *fr*, fasciculus retroflexus; *fx*, fornix; *LHA*, lateral hypothalamus; *LPO*, lateral preoptic area; *MRN*, midbrain reticular nucleus; *PAG*, periaqueductal gray; *pc*, posterior commissure; *PH*, posterior hypothalamus; *PVH*, paraventricular hypothalamic nucleus; *SC*, superior colliculus. Scale bars: A, E, G, I = 1mm; D, C = 500μm; B, D, F, H, J = 200μm.

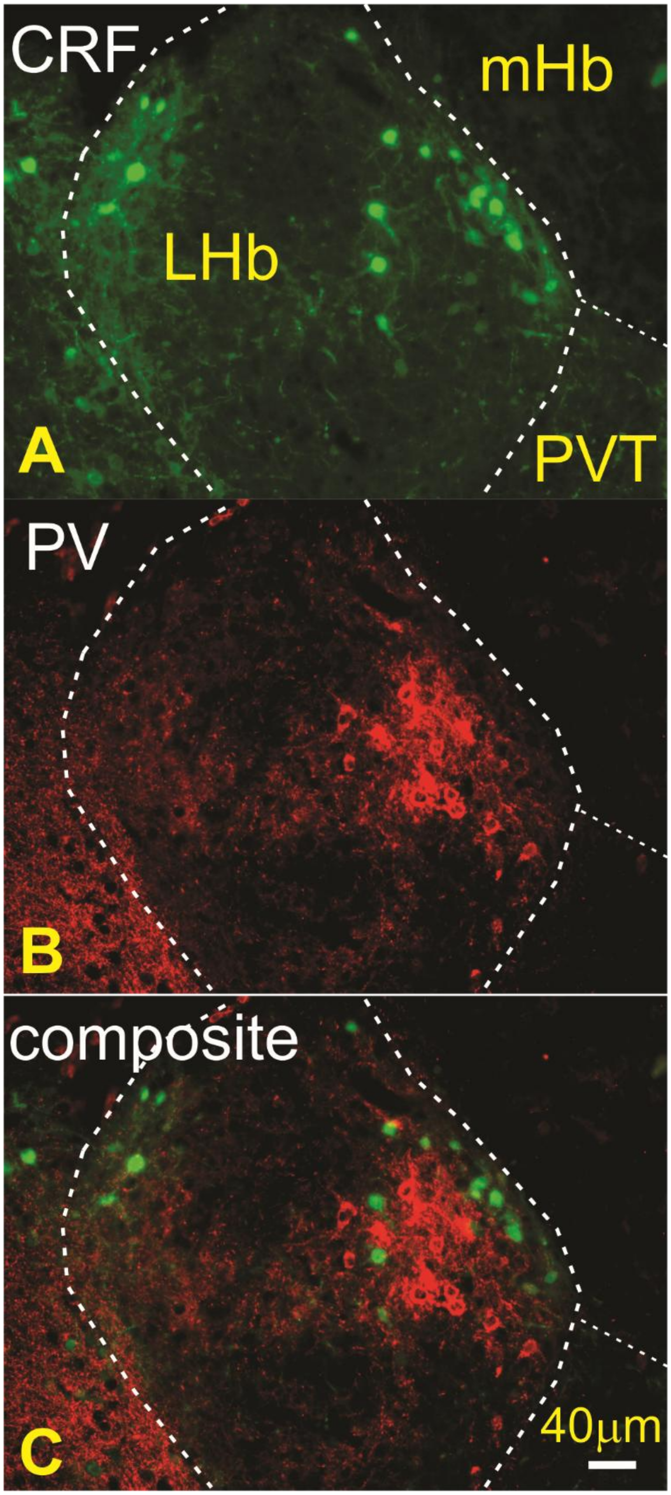
CRF
mHb
LHb
A
PVT
PV
B
composite
40μm
C

**Supplementary Figure 2: Neurochemical distinction between LHb CRF+ and PV+ Neurons**

Representative images of the mouse lateral habenula (LHb) characterizing the relationship between CRF-expressing and parvalbumin-expressing (PV+) neuronal populations.

- **(A)** LHb CRF+ Neurons: Visualization of retrogradely labeled CRF+ neurons (green) within the LHb.
- **(B)** Parvalbumin Immunostaining: Identification of PV+ neurons (red) using an anti-parvalbumin antibody.
- **(C)** Composite Image: Merged channels illustrating the spatial distribution of both populations.

Analysis of the composite image confirms that CRF+ and PV+ neurons constitute distinct, non-overlapping cell populations within the mouse LHb.

Abbreviations: *LHb*, lateral habenula; *PV*, parvalbumin.

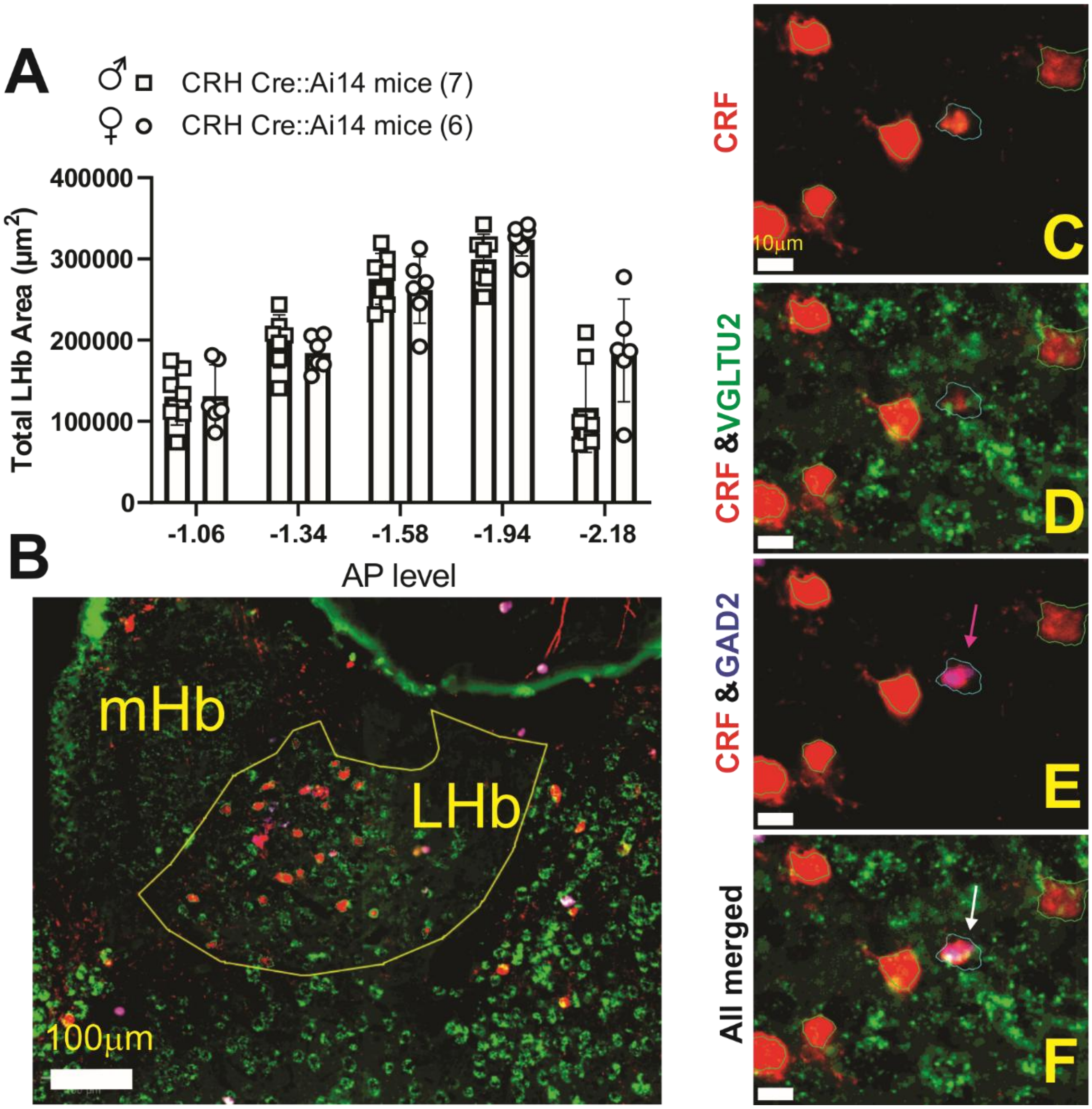


**Supplementary Figure 3: Quantitative Workflow and Anatomical Validation of LHb RNAscope Analysis**

**(A)** Volumetric Consistency across Sexes. Quantification of the total surface area (µm2) of the LHb across five anterior-posterior (AP) levels in male (squares, n=7) and female (circles, n=6) CRH-Cre::Ai14 mice. No significant sex differences were observed in the total area analyzed for the LHb perimeter, ensuring that subsequent cell count comparisons were not confounded by variations in regional volume. **(B)** QuPath Image Analysis Strategy. Representative low-magnification image illustrating the unbiased automated analysis workflow. The LHb was manually delineated as a Region of Interest (ROI), followed by the identification of CRF+ neurons (tdTomato) and the detection of multiplexed RNAscope transcripts.

**(C–F)** High-Resolution Phenotyping. Magnified representative images illustrating the subcellular localization of *Slc17a6* (vGLUT2) and *Gad2* mRNA within CRF+ neurons.

- **(C):** CRF+ (tdTomato) neurons alone.
- **(D):** Dual-channel overlay of CRF and *vGLUT2* (green), showing 100% colocalization.
- **(E):** Dual-channel overlay of CRF and *GAD2* (blue). The pink arrow highlights a discrete CRF+ neuron co-expressing *GAD2*.
- **(F):** All-channel merged image. The white arrow indicates the same *GAD2+/CRF+* neuron, further demonstrating its triple-positive status for *vGLUT2*.

Abbreviations: *LHb*, lateral habenula; *mHb*, medial habenula. Scale bars: B = 100 μm.; C-F = 10 μm.

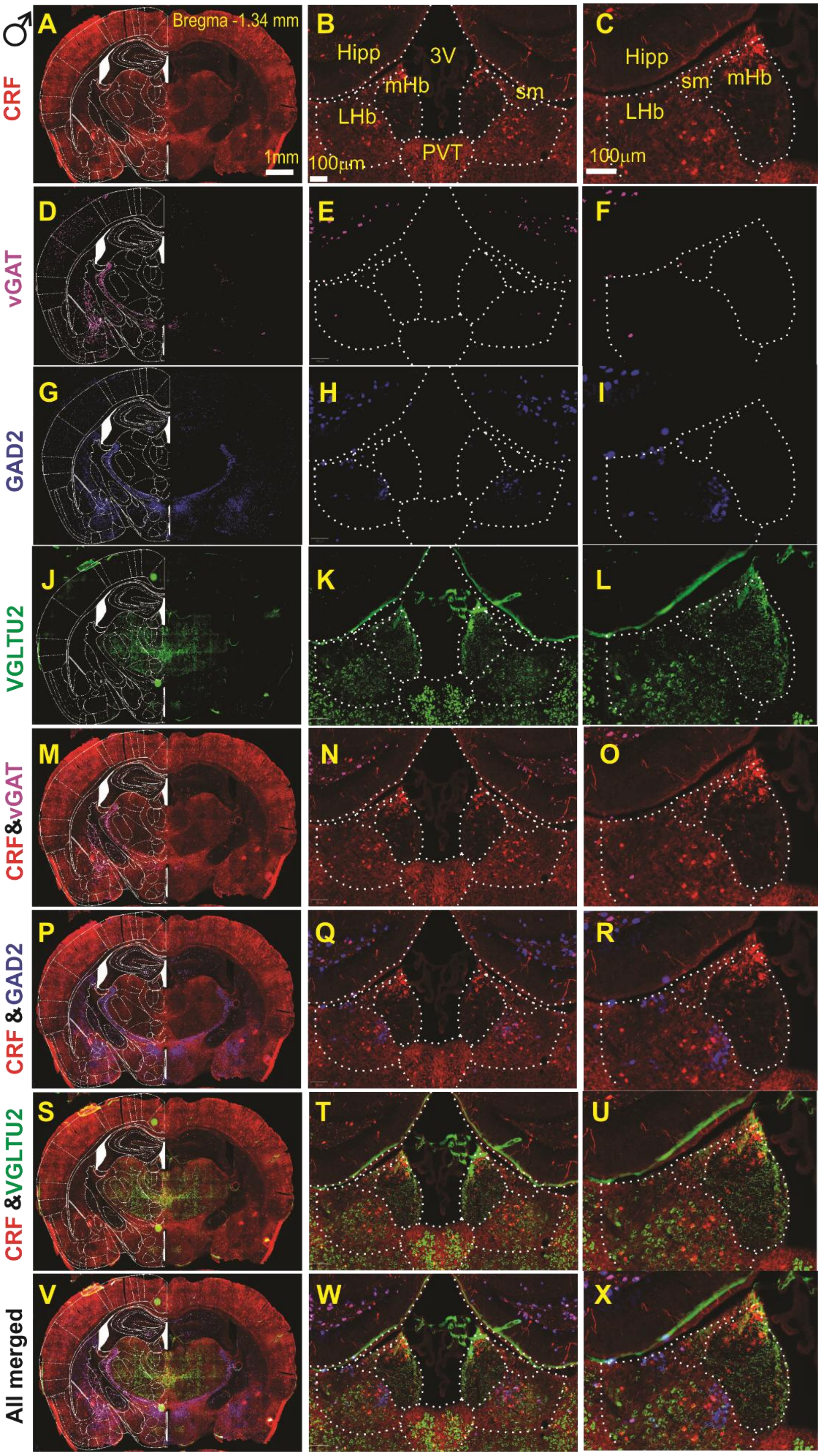

♂
CRF
vGAT
GAD2
VGLTU2
CRF&vGAT
CRF & GAD2
CRF &VGLTU2
All merged
A
Bregma -1.34 mm
1mm
B
Hipp
3V
mHb
sm
LHb
PVT
100μm
C
Hipp
sm
mHb
LHb
100μm
D
E
F
G
H
I
J
K
L
M
N
O
P
Q
R
S
T
U
V
W
X

**Supplementary Figure 4: Molecular Characterization of CRF+ Neurons in the male LHb at Bregma -1.34mm.**

Multiplexed RNAscope *in situ* hybridization was utilized to identify the neurotransmitter phenotype of CRF+ (tdTomato-labeled) neurons in male CRH-Cre::Ai14 mice at Bregma -1.34 mm. (**A**, **D**, **G**, **J**): Low-magnification hemicoronal views integrated with mouse atlas overlays, showing regional expression patterns of *Crh* (tdTomato), *Slc32a1* (vGAT), *Gad2* (GAD2), and *Slc17a6* (vGLUT2). (**B**–**L**): High-magnification images corresponding to panels A, D, G, and J, highlighting bilateral expression within the lateral habenula (LHb) and providing high-resolution details of the left LHb. (**M**–**U**): Dual-channel merged images demonstrating the colocalization (overlay) of CRF with inhibitory markers (*vGAT*, *GAD2*) and the excitatory marker (*vGLUT2*). (**V**–**X**): All-channel merged images illustrating the simultaneous colocalization of *vGAT*, *GAD2*, and *vGLUT2* within LHb CRF+ neurons. Abbreviations: *LHb*, lateral habenula; *mHb*, medial habenula; *Hipp*, hippocampus; *sm*, stria medullaris; *3V*, third ventricle; *PVT*, paraventricular nucleus of the thalamus. Scale bars: A = 1mm; B, C = 100 μm.

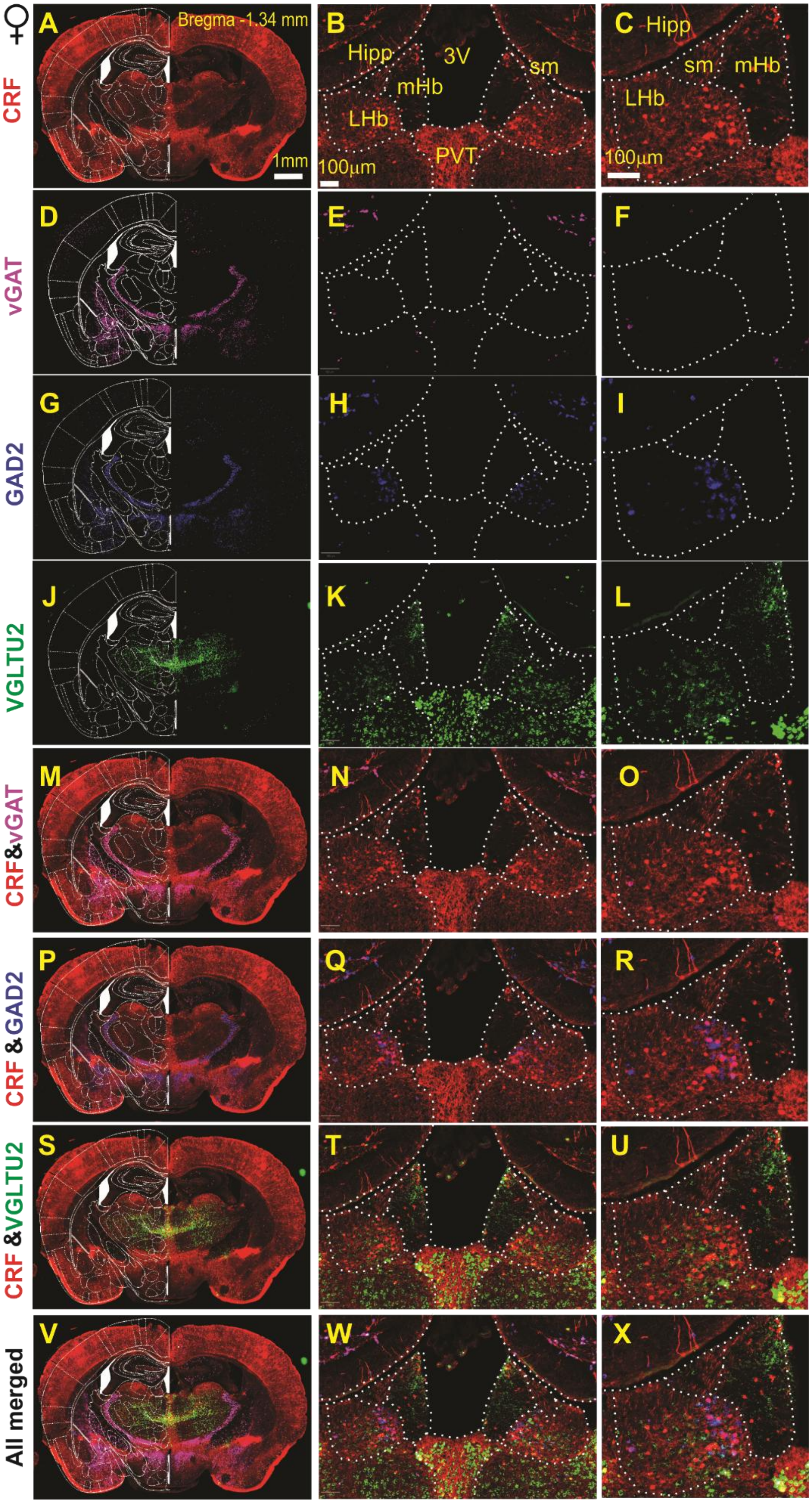

♀
CRF
vGAT
GAD2
VGLTU2
CRF&vGAT
CRF & GAD2
CRF &VGLTU2
All merged
A
Bregma -1.34 mm
1mm
B
Hipp
3V
sm
mHb
LHb
PVT
100µm
C
Hipp
sm
mHb
LHb
100µm
D
E
F
G
H
I
J
K
L
M
N
O
P
Q
R
S
T
U
V
W
X

**Supplementary Figure 5: Molecular Characterization of CRF+ Neurons in the female LHb at Bregma -1.34mm.**

Multiplexed RNAscope *in situ* hybridization was utilized to identify the neurotransmitter phenotype of CRF+ (tdTomato-labeled) neurons in female CRH-Cre::Ai14 mice at Bregma -1.34 mm. (**A**, **D**, **G**, **J**): Low-magnification hemicoronal views integrated with mouse atlas overlays, showing regional expression patterns of *Crh* (tdTomato), *Slc32a1* (vGAT), *Gad2* (GAD2), and *Slc17a6* (vGLUT2). (**B**–**L**): High-magnification images corresponding to panels A, D, G, and J, highlighting bilateral expression within the lateral habenula (LHb) and providing high-resolution details of the left LHb. (**M**–**U**): Dual-channel merged images demonstrating the colocalization (overlay) of CRF with inhibitory markers (*vGAT*, *GAD2*) and the excitatory marker (*vGLUT2*). (**V**–**X**): All-channel merged images illustrating the simultaneous colocalization of *vGAT*, *GAD2*, and *vGLUT2* within LHb CRF+ neurons. Abbreviations: *LHb*, lateral habenula; *mHb*, medial habenula; *Hipp*, hippocampus; *sm*, stria medullaris; *3V*, third ventricle; *PVT*, paraventricular nucleus of the thalamus. Scale bars: A = 1mm; B, C = 100 μm.

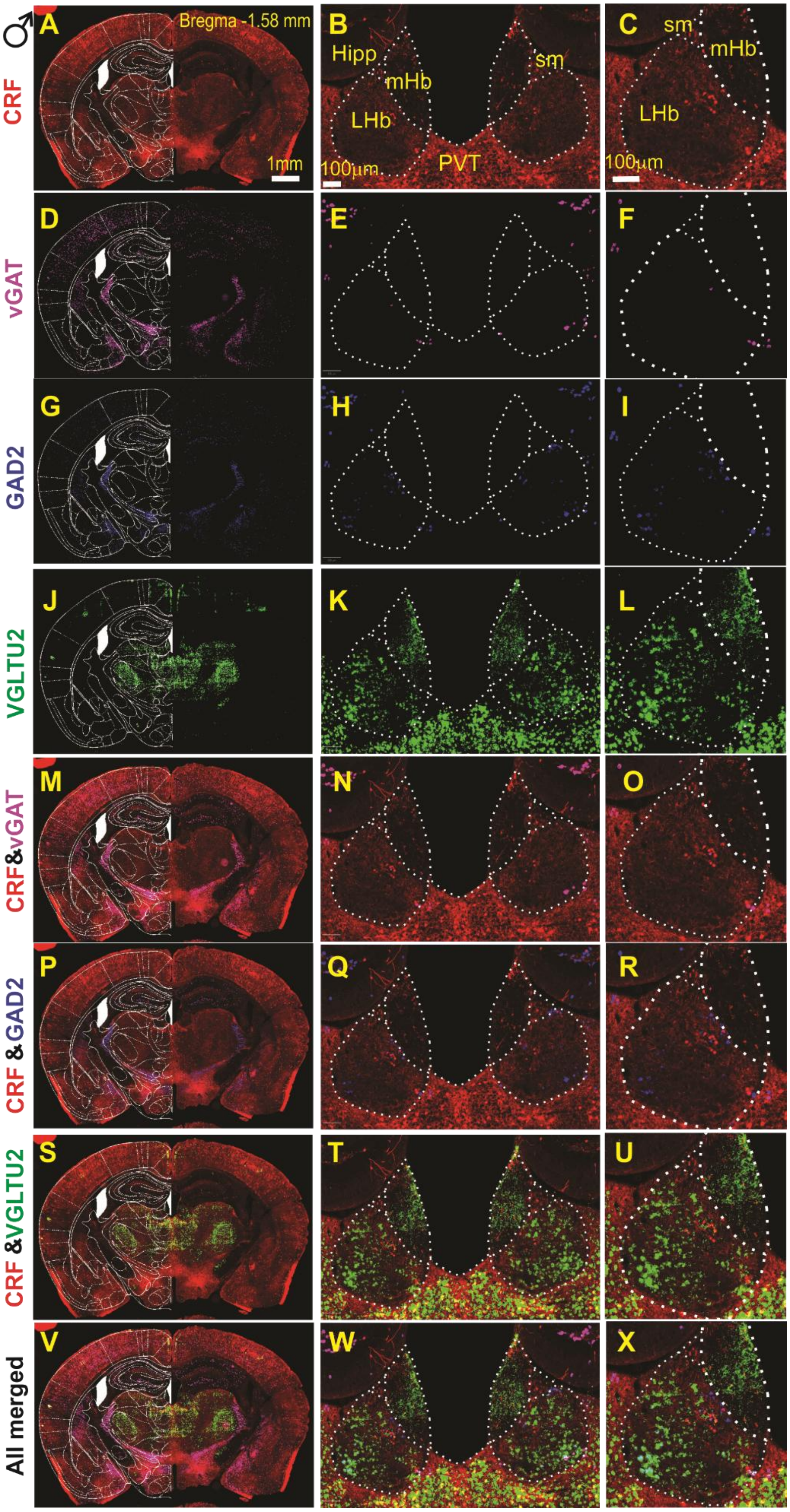
♂
CRF
vGAT
GAD2
VGLTU2
CRF&vGAT
CRF &GAD2
CRF &VGLTU2
All merged
A
Bregma -1.58 mm
1mm
B
Hipp
mHb
LHb
sm
PVT
100µm
C
sm
mHb
LHb
100µm
D
E
F
G
H
I
J
K
L
M
N
O
P
Q
R
S
T
U
V
W
X

**Supplementary Figure 6: Molecular characterization of CRF+ neurons in the male LHb at bregma -1.58mm.**

Multiplexed RNAscope *in situ* hybridization was utilized to identify the neurotransmitter phenotype of CRF+ (tdTomato-labeled) neurons in male CRH-Cre::Ai14 mice at Bregma -1.58 mm. (**A**, **D**, **G**, **J**): Low-magnification hemicoronal views integrated with mouse atlas overlays, showing regional expression patterns of *Crh* (tdTomato), *Slc32a1* (vGAT), *Gad2* (GAD2), and *Slc17a6* (vGLUT2). (**B**–**L**): High-magnification images corresponding to panels A, D, G, and J, highlighting bilateral expression within the lateral habenula (LHb) and providing high-resolution details of the left LHb. (**M**–**U**): Dual-channel merged images demonstrating the colocalization (overlay) of CRF with inhibitory markers (*vGAT*, *GAD2*) and the excitatory marker (*vGLUT2*). (**V**–**X**): All-channel merged images illustrating the simultaneous colocalization of *vGAT*, *GAD2*, and *vGLUT2* within LHb CRF+ neurons. Abbreviations: *LHb*, lateral habenula; *mHb*, medial habenula; *Hipp*, hippocampus; *sm*, stria medullaris; *3V*, third ventricle; *PVT*, paraventricular nucleus of the thalamus. Scale bars: A = 1mm; B, C = 100 μm.

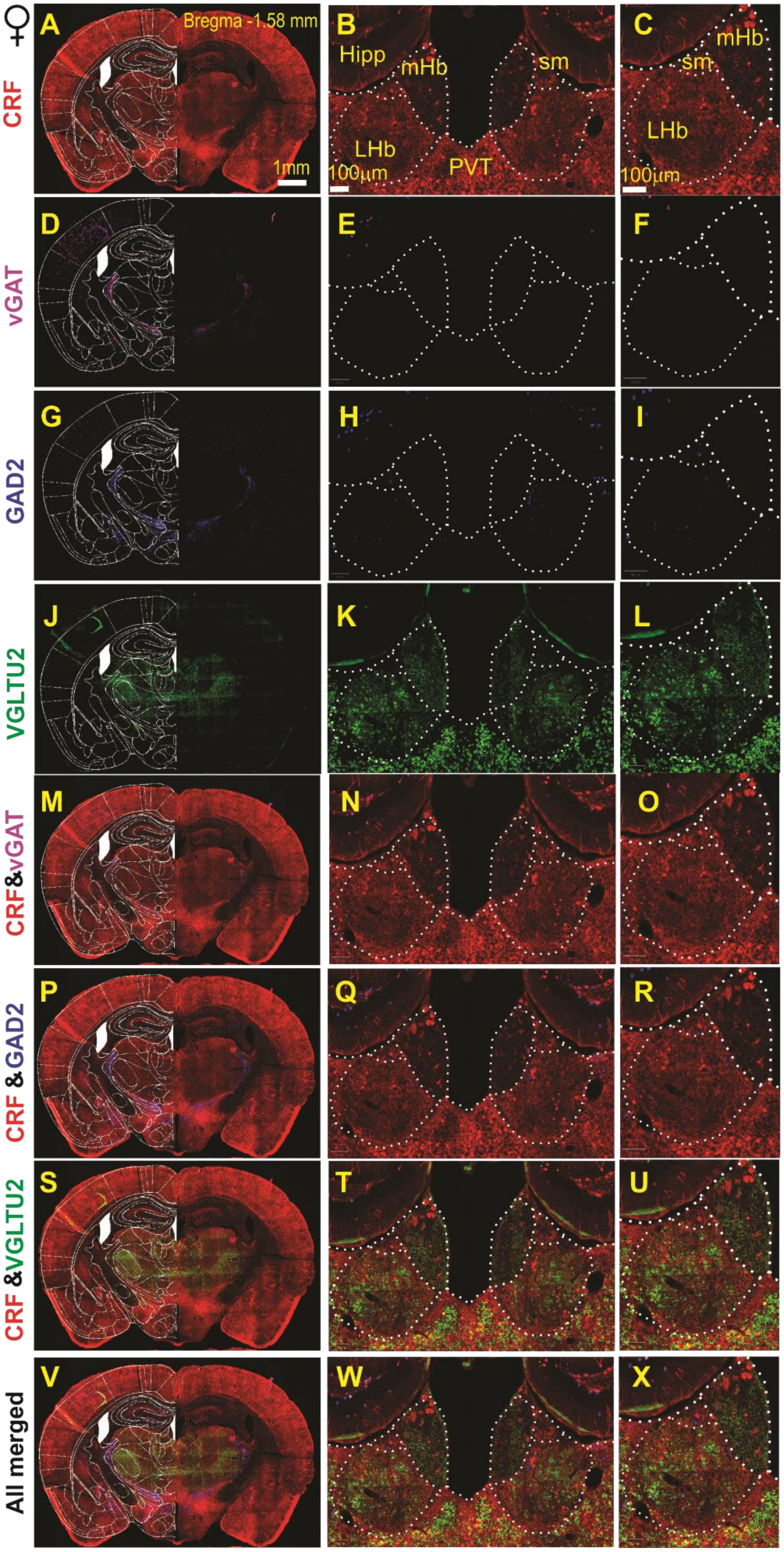

♀
CRF
vGAT
GAD2
VGLTU2
CRF&vGAT
CRF & GAD2
CRF &VGLTU2
All merged
A
Bregma -1.58 mm
1mm
B
Hipp
mHb
sm
LHb
PVT
100μm
C
mHb
sm
LHb
100μm
D
E
F
G
H
I
J
K
L
M
N
O
P
Q
R
S
T
U
V
W
X

**Supplemental Figure 7: Molecular characterization of CRF+ neurons in the female LHb at bregma -1.58mm.**

Multiplexed RNAscope *in situ* hybridization was utilized to identify the neurotransmitter phenotype of CRF+ (tdTomato-labeled) neurons in female CRH-Cre::Ai14 mice at Bregma -1.58 mm. (**A**, **D**, **G**, **J**): Low-magnification hemicoronal views integrated with mouse atlas overlays, showing regional expression patterns of *Crh* (tdTomato), *Slc32a1* (vGAT), *Gad2* (GAD2), and *Slc17a6* (vGLUT2). (**B**–**L**): High-magnification images corresponding to panels A, D, G, and J, highlighting bilateral expression within the lateral habenula (LHb) and providing high-resolution details of the left LHb. (**M**–**U**): Dual-channel merged images demonstrating the colocalization (overlay) of CRF with inhibitory markers (*vGAT*, *GAD2*) and the excitatory marker (*vGLUT2*). (**V**–**X**): All-channel merged images illustrating the simultaneous colocalization of *vGAT*, *GAD2*, and *vGLUT2* within LHb CRF+ neurons. Abbreviations: *LHb*, lateral habenula; *mHb*, medial habenula; *Hipp*, hippocampus; *sm*, stria medullaris; *3V*, third ventricle; *PVT*, paraventricular nucleus of the thalamus. Scale bars: A = 1mm; B, C = 100 μm.

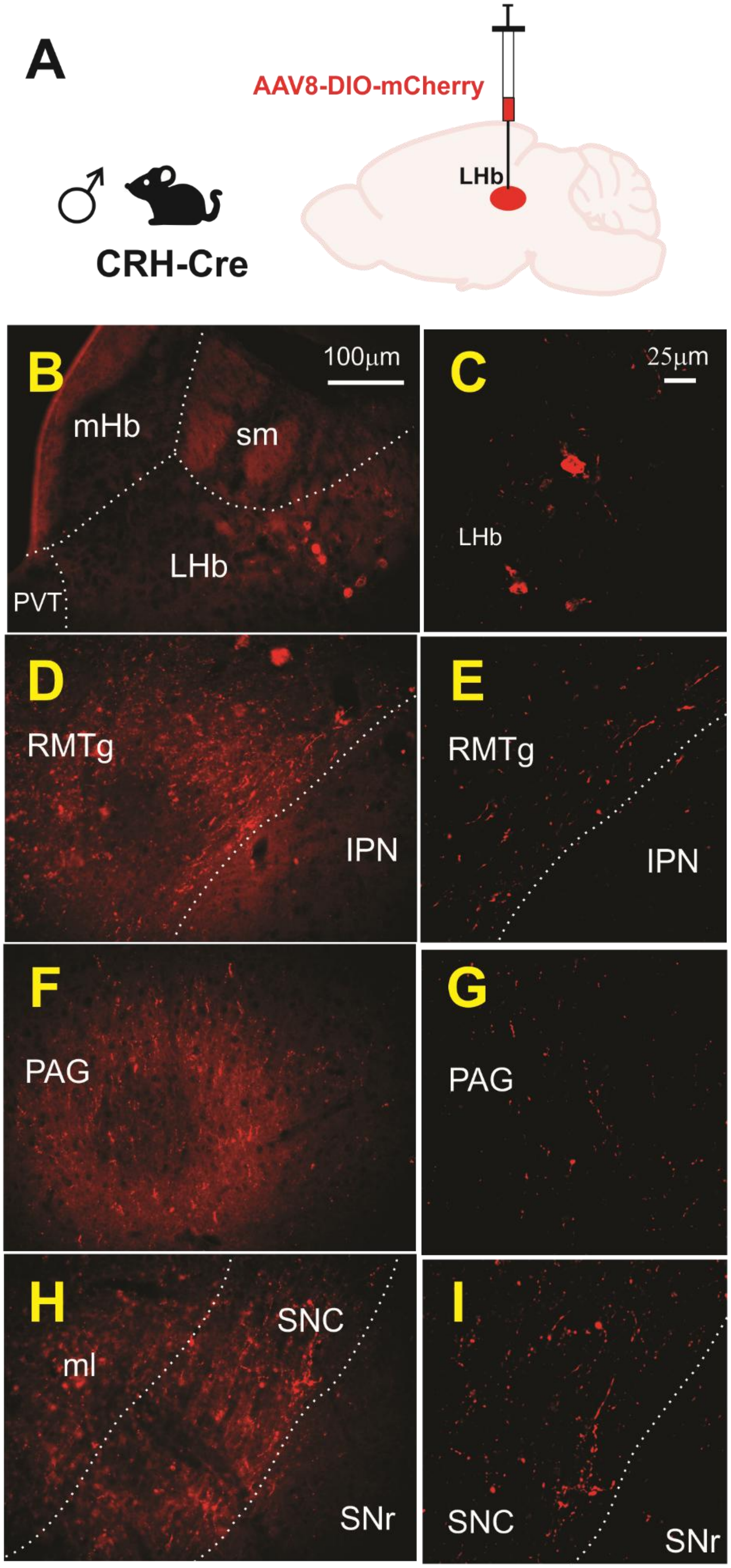
A
AAV8-DIO-mCherry
LHb
♂
CRH-Cre
B
100μm
mHb
sm
LHb
PVT
C
25μm
LHb
D
RMTg
IPN
E
RMTg
IPN
F
PAG
G
PAG
H
SNC
ml
SNr
I
SNC
SNr

**Supplementary Figure 8: Anterograde mapping of $LHb^{CRF}$ projections using mCherry in male mice**

**(A)** Schematic of the viral approach: a Cre-dependent anterograde vector (AAV8-DIO-mCherry) was injected into the LHb of male CRH-Cre mice (n=3).

**(B–I)** Representative Projection Targets. Fluorescent microscopy images showing the distribution of mCherry-labeled axonal processes originating from LHb CRF+ neurons. Panels on the left illustrate lower magnification overviews, while panels on the right provide corresponding high-magnification details of labeled fibers.

- **(B, C):** Local innervation within the LHb and surrounding mHb.
- **(D, E):** Projections to the rostromedial tegmental nucleus (RMTg) and adjacent interpeduncular nucleus (IPN).
- **(F, G):** Axonal fibers within the periaqueductal gray (PAG).
- **(H, I):** Innervation of the substantia nigra pars compacta (SNC) and neighboring substantia nigra pars reticulata (SNr).

Abbreviations: *IPN*, interpeduncular nucleus; *mHb*, medial habenula; *ml*, medial lemniscus; *PVT*, paraventricular nucleus of the thalamus; *sm*, stria medullaris; *SNC*, substantia nigra pars compacta; *SNr*, substantia nigra pars reticulata. Scale bars: B = 100 μm; C = 25 μm.

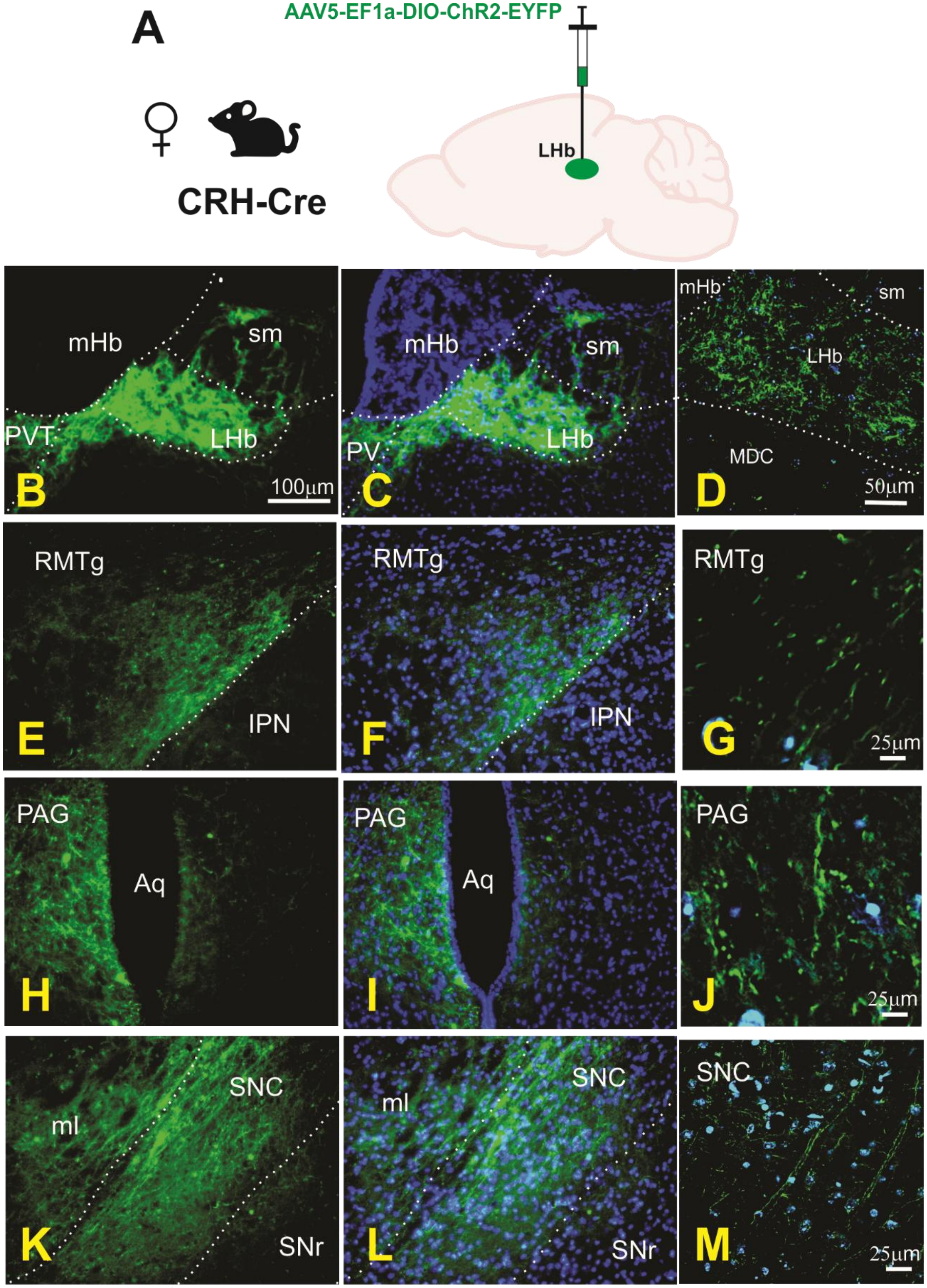

A
AAV5-EF1a-DIO-ChR2-EYFP
LHb
CRH-Cre
mHb
sm
PVT
LHb
B
100µm
mHb
sm
PV
LHb
C
mHb
sm
LHb
MDC
D
50µm
RMTg
IPN
E
RMTg
IPN
F
RMTg
G
25µm
PAG
Aq
H
PAG
Aq
I
PAG
J
25µm
ml
SNC
SNr
K
ml
SNC
SNr
L
SNC
M
25µm

**Supplementary Figure 9: Anterograde mapping of LHb$^{CRF}$ projections using ChR2-EYFP in female mice**

**(A)** Schematic of the viral approach: a Cre-dependent anterograde vector (AAV5-EF1a-DIO-ChR2-EYFP) was injected into the LHb of female CRH-Cre mice (n=3). **(B–M)** Fluorescent and confocal microscopy images showing EYFP-labeled axonal processes originating from LHb CRF+ neurons. Panels **B**, **E**, **H**, and **K** show overviews of EYFP expression; panels **C**, **F**, **I**, and **L** show the same overviews integrated with DAPI (blue) nuclear staining; and panels **D**, **G**, **J**, and **M** provide high-magnification details of labeled fibers.

- **(B–D):** Local terminal innervation within the LHb and surrounding mHb.
- **(E–G):** Projections to the rostromedial tegmental nucleus (RMTg) and adjacent IPN.
- **(H–J):** Axonal fibers within the periaqueductal gray (PAG) flanking the aqueduct (Aq).
- **(K–M):** Dense innervation of the substantia nigra pars compacta (SNC) near the medial lemniscus (ml) and substantia nigra pars reticulata (SNr).

Abbreviations: *Aq*, aqueduct; *IPN*, interpeduncular nucleus; *mHb*, medial habenula; *ml*, medial lemniscus; *MDC*, mediodorsal nucleus of the thalamus; *PVT*, paraventricular nucleus of the thalamus; *sm*, stria medullaris; *SNC*, substantia nigra pars compacta; *SNr*, substantia nigra pars reticulata. Scale bars: B, E, H, K = 100 μm; D = 50 μm; G, J, M = 25 μm.

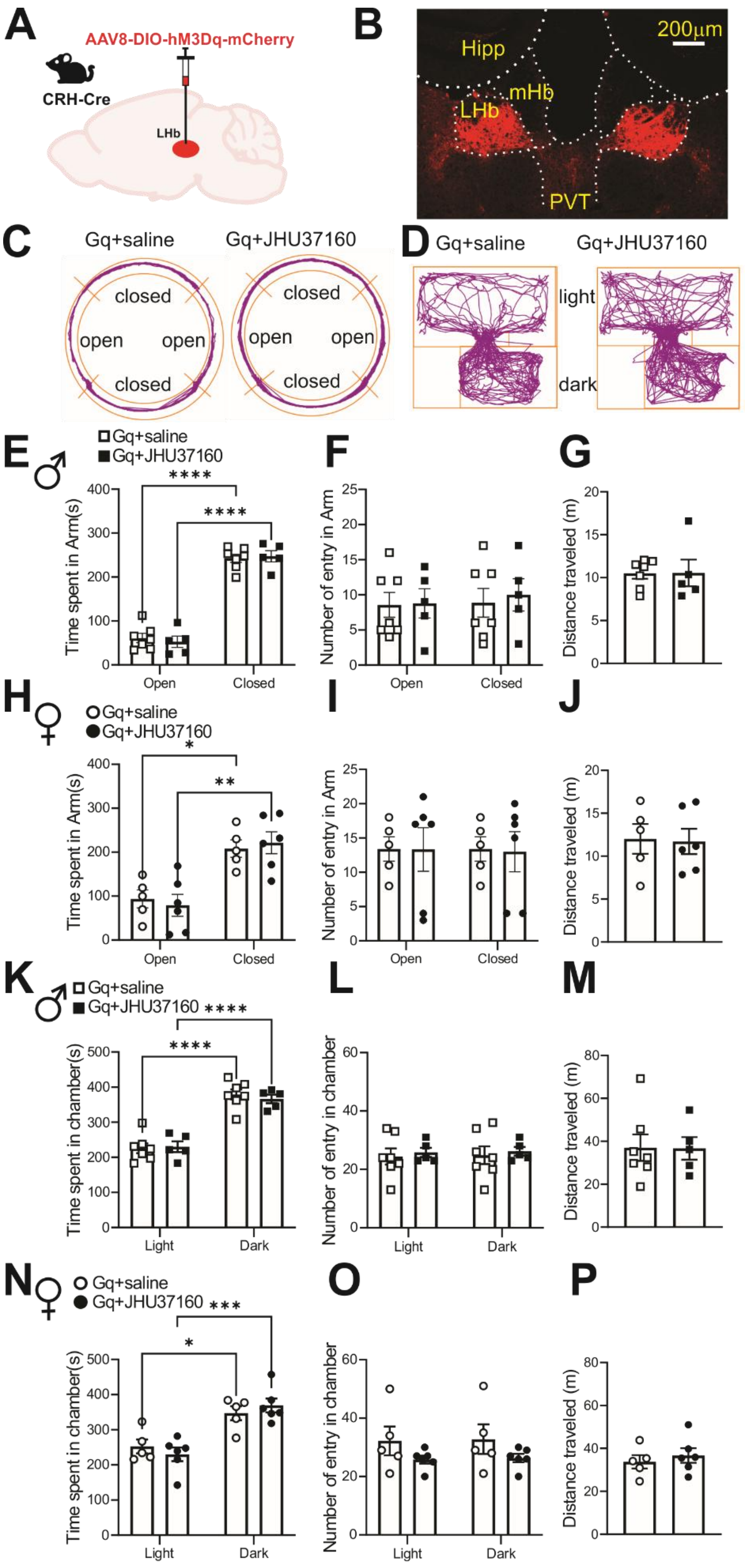

A
AAV8-DIO-hM3Dq-mCherry
CRH-Cre
LHb
B
200μm
Hipp
mHb
LHb
PVT
C
Gq+saline
Gq+JHU37160
closed
open
open
closed
D
Gq+saline
Gq+JHU37160
light
dark
E ♂
Gq+saline
Gq+JHU37160
Time spent in Arm(s)
Open
Closed
F
Number of entry in Arm
G
Distance traveled (m)
H ♀
I
J
K ♂
Time spent in chamber(s)
Light
Dark
L
Number of entry in chamber
M
N ♀
O
P

**Supplementary Figure 10: Chemogenetic activation of $LHb^{CRF}$ neurons did not affect locomotor activity or anxiety-related behavior in the elevated zero maze (EZM) or light-dark box (LDT). A** Schematic depicting AAV8-DIO-hM3Dq-mCherry injection into the LHb of a CRH-Cre mouse. **B:** Representative coronal section verifying the restricted expression of hM3Dq-mCherry within $LHb^{CRF}$ neurons. **C-D** Representative track plots illustrating the movement of mice in the EZM (**C**) and LDT (**D**) following saline or JHU37160 injection. **E-J** Quantification of EZM behavior, showing no difference between JHU37160- and saline-injected male (**E–G**; squares) or female (**H–J**; circles) mice in time spent in open arms, number of entries to open arms, or total distance traveled. **K-P** Quantification of LDT behavior, showing no difference between JHU37160- and saline-injected male **(K–M**; squares**)** or female **(N–P**; circles**)** mice in time spent in light chambers, number of entries to light chambers, or total distance traveled. Abbreviations: *Hipp*, hippocampus; *mHb*, medial habenula; *PVT*, paraventricular nucleus of the thalamus. *p<0.05, **p<0.01, ***p<0.001, ****p<0.0001; 2way-ANOVA.
